# Discovery of multiple anti-CRISPRs uncovers anti-defense gene clustering in mobile genetic elements

**DOI:** 10.1101/2020.05.22.110304

**Authors:** Rafael Pinilla-Redondo, Saadlee Shehreen, Nicole D. Marino, Robert D. Fagerlund, Chris M. Brown, Søren J. Sørensen, Peter C. Fineran, Joseph Bondy-Denomy

## Abstract

Many prokaryotes employ CRISPR-Cas systems to combat invading mobile genetic elements (MGEs). In response, some MGEs have evolved Anti-CRISPR (Acr) proteins to bypass this immunity, yet the diversity, distribution and spectrum of activity of this immune evasion strategy remain largely unknown. Here, we uncover 11 new type I anti-CRISPR genes encoded on numerous chromosomal and extrachromosomal mobile genetic elements within *Enterobacteriaceae* and *Pseudomonas*. Candidate genes were identified adjacent to anti-CRISPR associated gene 5 (*aca5)* and assayed against a panel of six type I systems: I-F (*Pseudomonas*, *Pectobacterium*, and *Serratia*), I-E (*Pseudomonas* and *Serratia*), and I-C (*Pseudomonas)*, revealing the type I-F and/or I-E *acr* genes and a new *aca* (*aca9*). We find that *acr* genes not only associate with other *acr* genes, but also with inhibitors of distinct bacterial defense systems. These genomic regions appear to be “anti-defense islands”, reminiscent of the clustered arrangement of “defense islands” in prokaryotic genomes. Our findings expand on the diversity of CRISPR-Cas inhibitors and reveal the potential exploitation of *acr* loci neighborhoods for identifying new anti-defense systems.

## Introduction

All cellular life is under the constant threat of invasion by foreign genetic elements. In particular, prokaryotes are outnumbered by a wide spectrum of mobile genetic elements (MGEs) that infect them, including viruses and plasmids. This selective pressure has driven the evolution of diverse defense mechanisms (Rostøl and Marraffini, 2019; Hampton, Watson and Fineran, 2020), including clustered regularly interspaced short palindromic repeats (CRISPR) and CRISPR-associated (Cas) genes, a family of adaptive immune systems.

CRISPR-Cas loci have been identified in sequenced genomes of around 40% of bacteria and 85% of archaea (Makarova *et al.*, 2020), bearing testament to their evolutionary and ecological importance. This mode of defense allows cells to remember, recognize and thwart recurrently infecting agents. Broadly, CRISPR-Cas immunity consists of three main stages: adaptation, processing/biogenesis, and interference (Hille *et al.*, 2018). During adaptation, snippets of an invading genetic element are incorporated into CRISPR arrays as “spacers” between repeat sequences, yielding a heritable record of former genetic intruders. The CRISPR array is then expressed as a long transcript (pre-crRNA) that is processed into single CRISPR RNAs (crRNAs), which guide Cas nucleases to target invading nucleic acids that carry a complementary sequence to the spacer (referred to as protospacer).

In response to the strong selective pressure exerted by CRISPR-Cas immunity, many MGEs have developed inhibitors of CRISPR-Cas function called anti-CRISPR (Acr) proteins (Borges, Davidson and Bondy-Denomy, 2018). The first *acr* genes were discovered in phages that inhibit the type I-F CRISPR-Cas system of *Pseudomonas aeruginosa* (Bondy-Denomy *et al.*, 2013). Many more non-homologous Acr proteins have been subsequently reported for different CRISPR-Cas types (e.g. types II, III and V) (Pawluk, Amrani, *et al.*, 2016; Rauch *et al.*, 2017; Marino *et al.*, 2018; Watters *et al.*, 2018; Bhoobalan-Chitty *et al.*, 2019; Athukoralage *et al.*, 2020), including some on non-phage MGEs (Mahendra *et al.*, 2020). Previous structural and biochemical characterization of Acr proteins has revealed a diverse range of inhibitory activities, including interference with crRNA loading, inhibition of target DNA recognition, and inhibition of DNA cleavage, among others (Davidson *et al.*, 2020). Apart from sharing a typically low molecular weight, Acrs lack conserved sequence and structural features, thus rendering *de novo* prediction largely impractical with current methods. However, *acr* genes tend to cluster within loci that encode more conserved Acr-associated (Aca) proteins (Pawluk, Staals, *et al.*, 2016; Borges, Davidson and Bondy-Denomy, 2018), which are transcriptional repressors of the *acr* locus (Birkholz *et al.*, 2019; Stanley *et al.*, 2019). The *aca* genes are often more broadly distributed than *acr* genes and have been used to uncover new *acr* loci (Pawluk, Amrani, *et al.*, 2016; Pawluk, Staals, *et al.*, 2016; Lee *et al.*, 2018; Marino *et al.*, 2018).

The discovery of Acr proteins explains how MGEs can persist despite frequent targeting by host spacer sequences. Acrs are therefore a strong driver for the diversification of CRISPR-Cas systems in nature and possibly the accumulation of other defense systems in prokaryotic genomes. Because MGEs facilitate host genome rearrangements and provide the foundation for vast prokaryotic gene exchange networks, the study of Acrs allows a better understanding of MGE-host interactions and the horizontal transfer potential of MGE-encoded traits (e.g. antibiotic resistance) across microbiomes. Practically, Acr proteins also benefit phage-based therapeutics and plasmid-based delivery platforms and provide a means to control CRISPR-Cas-derived biotechnologies (Marino *et al.*, 2020).

In this study, we investigate the interactions between MGEs and their bacterial hosts, focusing on uncovering new Acrs that enable MGEs to avoid potent host defense mechanisms. We describe the discovery of 11 type I-F and/or I-E Acr families encoded by phage and non-phage MGEs by leveraging *aca5* and a newly identified *aca* gene (*aca9*) as markers for *acr* loci. Bioinformatic analyses further revealed that *acrs* cluster with other anti-defense systems within MGE genomes, suggesting the existence of “anti-defense islands” and highlighting a potential avenue for the discovery of unknown anti-defense genes.

## Results

### An *aca5*-based computational search for Acr candidates

To uncover how MGEs within the Enterobacteriales order cope with the pressure of CRISPR-Cas immunity, we performed bioinformatic searches using the Enterobacteriales-enriched *aca5* gene (Marino *et al.*, 2018). These searches revealed a wide phylogenetic distribution of homologs across bacterial families (Figure 1a), including members of the *Salmonella, Pectobacterium, Klebsiella, Serratia*, and *Escherichia* genera. Distant homologs were also identified in other bacterial orders (e.g. Vibrionales) at a considerably lower prevalence. Importantly, these organisms are enriched with class 1 CRISPR-Cas systems (primarily types I-F and/or I-E) (Makarova *et al.*, 2020), suggesting that their MGEs may rely on unknown type I inhibitors to bypass immunity.

**Figure 1.**
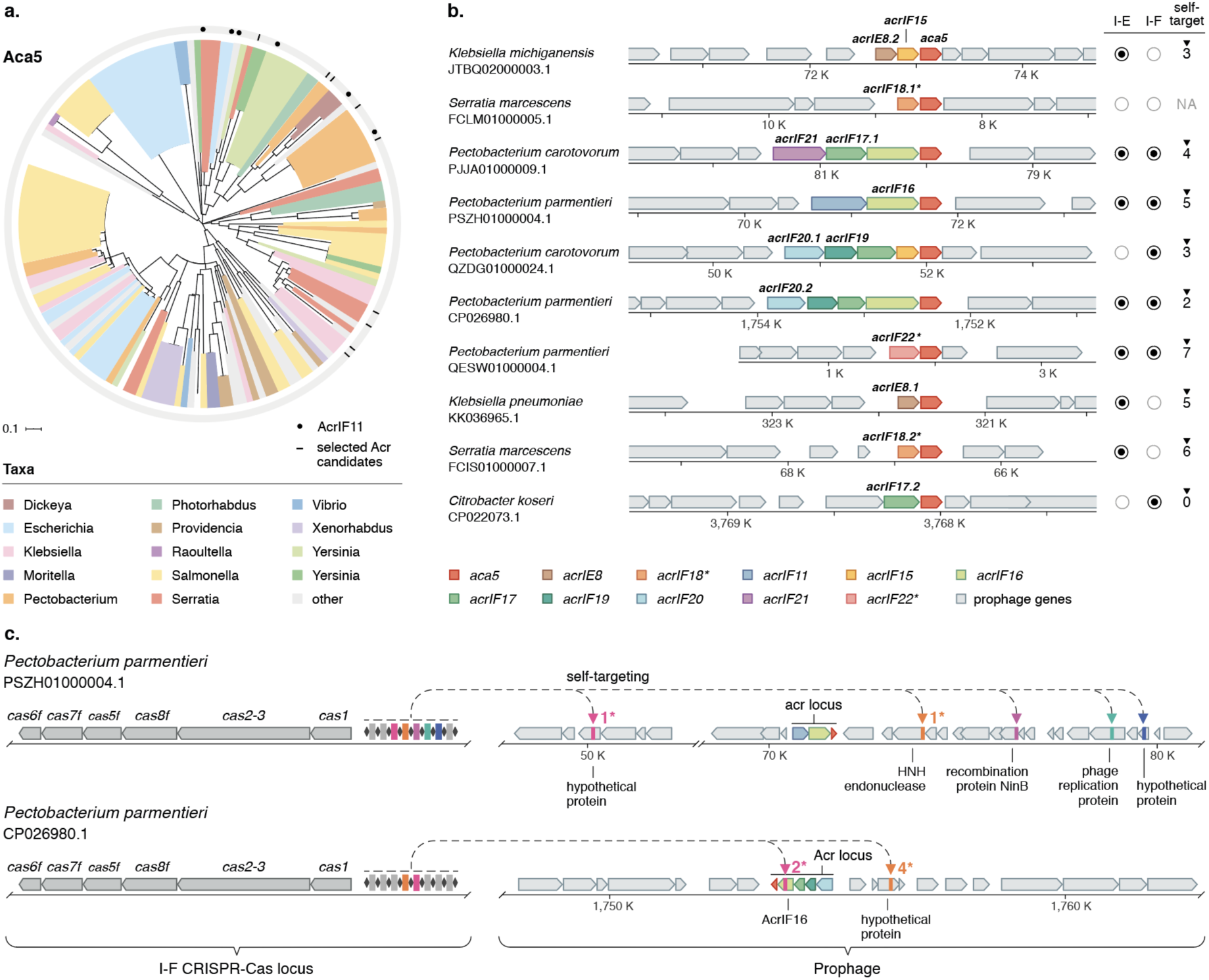
Bioinformatic search of *aca5*-associated *acrs*. **(a)** Phylogenetic distribution of Aca5 homologs across bacterial taxa. Tree branches are color-coded according to the genus from which the Aca5 orthologs originate. Black circles in the outer ring depict instances where Aca5 is associated with AcrIF11, marking the starting points for the guilt-by-association search; black lines represent the genomes from which *acr* candidates were selected for functional testing. **(b)** Genomic organization of the *acrs* selected for testing. Genes are colored by Acr family and the surrounding prophage genomic contexts are depicted in gray. Specific orthologs selected for testing are highlighted in bold. The presence/absence of I-F and I-E CRISPR-Cas systems and self-targeting spacers in the host genomes are summarized on the right (for details see Supplementary Datasheets S1 and S2). Self-targeting analysis was not possible in genomes where CRISPR-Cas systems were not detected, marked here as “NA”. **(c)** Graphic representation of the self-targeting instances observed in two *Pectobacterium parmentieri* genomes from which *acrIF16* and *acrIF20* genes were selected for testing. Colored arrowheads indicate the positions within the prophages that are targeted by a spacer in one of the host’s I-F CRISPR arrays. Genes are colored according to the legend in panel “b”. When possible, the name of the targeted gene is provided. Asterisks next to arrowheads indicate the number of spacer-protospacer mismatches.

Based on common characteristics of known Acr proteins, we restricted our candidate list to small predicted proteins (<200 amino acids) encoded upstream of *aca5* within genomic regions containing numerous MGE-associated genes. Following this approach, we identified several *acr* candidates residing in prophages from genomes of *Pectobacterium*, *Serratia*, *Klebsiella* and *Citrobacter*, and 10 genomes harboring a diverse set of putative *acr* genes were selected for further study (Figure 1b, Supplementary Datasheet S1). Notably, while the position of *aca5* remained fixed across these putative *acr* operons, *acr* candidates often co-occurred in shuffled clusters of 2-4 genes, as seen for other *acr* loci (Bondy-Denomy *et al.*, 2013; Pawluk *et al.*, 2014; Rauch *et al.*, 2017; Marino *et al.*, 2018).

Bacteria that express *acrs* can tolerate “self-targeting” spacers that, in the absence of CRISPR-Cas inhibition, would otherwise cause lethal genomic cleavage (Bondy-Denomy *et al.*, 2013; Vercoe *et al.*, 2013; Xu *et al.*, 2019; Rollie *et al.*, 2020). The presence of these self-targeting spacers can therefore be used to identify bacterial genomes that likely encode anti-CRISPR proteins capable of inhibiting their endogenous CRISPR system (Rauch *et al.*, 2017; Marino *et al.*, 2018; Watters *et al.*, 2018, 2020). We found that 8 out of the 10 selected genomes contained several I-F and/or I-E self-targeting spacers (Figure 1b, right, Supplementary Datasheets S1 and S2). A large number of these spacers (21/34 - 62% of all self-targeting hits) matched targets within the predicted prophages carrying the *acr* candidates (Supplementary Datasheet S2). Due to the promiscuous PAM of I-E (Westra *et al.*, 2013; Fineran *et al.*, 2014; Gleditzsch *et al.*, 2019), we were unable to confidently ascertain whether the PAM would enable targeting of the predicted spacer-protospacer matches. However, most I-F protospacers (83%) were flanked by the conserved I-F 5’-GG-3’ PAM, as described previously (Cady *et al.*, 2012; Rollins *et al.*, 2015) (Figure 1c, Supplementary Datasheet S2). Collectively, we concluded that the identified prophage genomes likely harbored *acr* loci and proceeded to experimentally test the 9 selected type I-E/I-F candidate *acr* genes.

### Newly identified Acr proteins inhibit different type I CRISPR-Cas systems

The nine candidate *acr* genes (and 4 additional homologs, denoted here by a decimal number after the Acr name) were tested against a panel of type I CRISPR-Cas systems: two type I-E variants (*Serratia* sp. ATCC39006 and *Pseudomonas aeruginosa* SMC4386), three type I-F variants (*Serratia* sp. ATCC39006, *Pseudomonas aeruginosa* PA14, and *Pectobacterium atrosepticum* SCRI1043), and one type I-C system (engineered strain of *Pseudomonas aeruginosa* PAO1). Notably, the different CRISPR-Cas subtypes and variants in the chosen model organisms span a wide phylogenetic diversity. While some of the tested systems display high degrees of similarity (e.g. >80% aa identity) with the CRISPR-Cas systems present in the endogenous *acr* hosts, others exhibit high divergence (<50% aa identity) (Figure 2, Supplementary Datasheet S3), allowing us to test for possible differences in the breadth of inhibitory activity. We challenged each of the model organisms with a CRISPR-targeted phage and assayed whether its replication could be restored by an *acr* candidate gene (Figure 2a).

**Figure 2.**
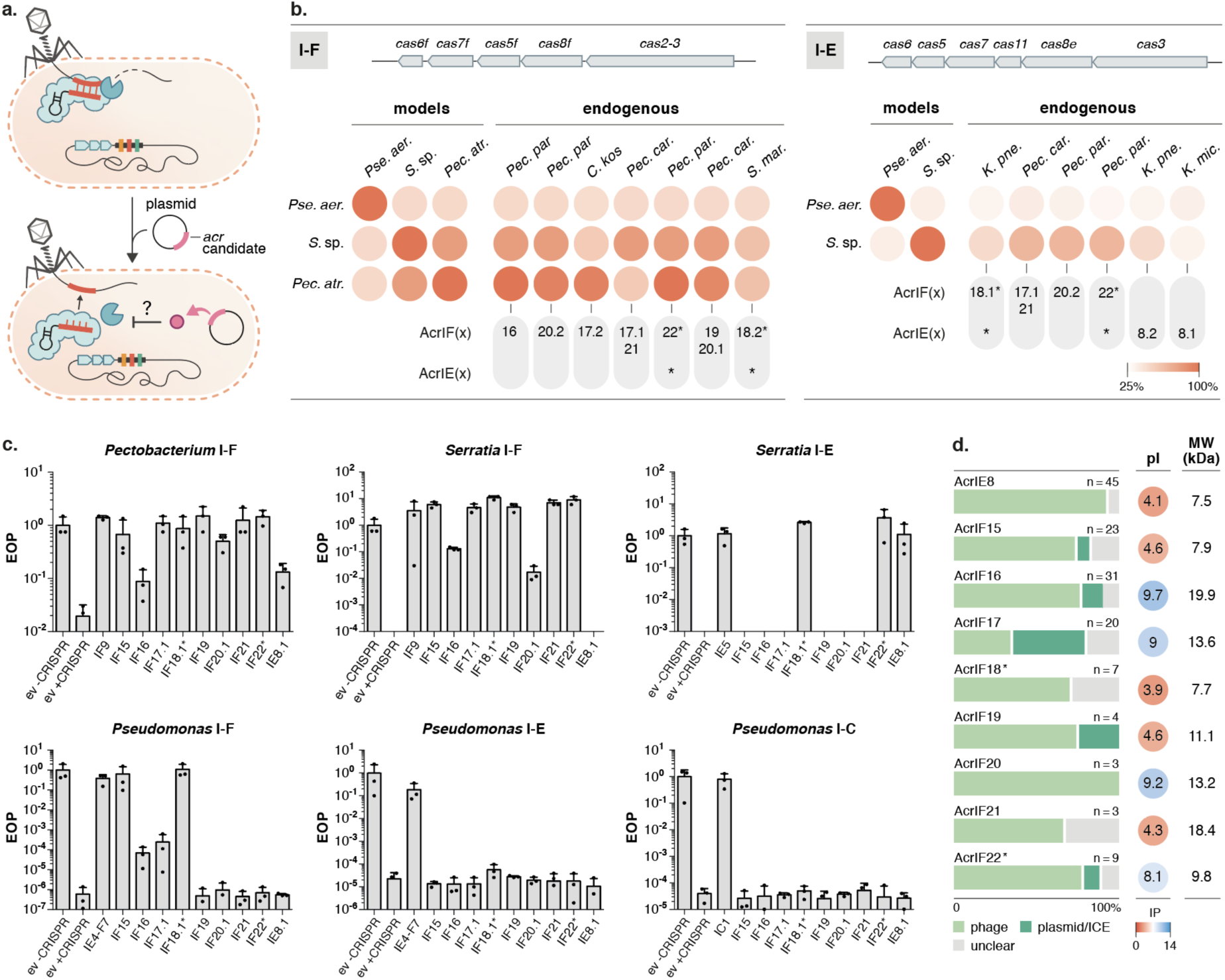
Newly identified *acr* genes inhibit diverse type I CRISPR-Cas systems. **(a)** Schematic of the experimental setup employed for *acr* validation. Bacteria carrying their native type I-F, I-E, or I-C CRISPR-Cas systems and a plasmid expressing each *acr* candidate were challenged with a CRISPR-targeted phage and infectivity was assessed. **(b)** Percent sequence identity comparison between the model CRISPR-Cas systems challenged (type I-F: *Pectobacterium*, *Serratia*, and *Pseudomonas*; type I-E: *Serratia* and *Pseudomonas;* type I-C: *Pseudomonas*) and the type I-F/I-E systems found in the endogenous hosts of the *acr* candidates, when available. Average values for the individual percentage identity comparisons between Cas orthologs forming the type I-E/I-F interference complexes are shown, excluding the adaptation modules (see top gene map) (Supplementary Datasheet S3). The specific Acr orthologs found in the genomes which were selected for testing are noted below; asterisks indicate dual anti-I-F/I-E function. **(c)** Efficiency of plaquing (EOP) of CRISPR-targeted phages in bacterial lawns expressing the different *acr* candidates, or the empty vector control (ev +CRISPR), compared to EOP of the same phage in non-targeting bacterial lawns carrying the empty vector (ev -CRISPR). Positive controls for Acr inhibition include AcrIF9 (*Pectobacterium* and *Serratia* I-F), AcrIE5 (*Serratia* I-E), AcrIE4-F7 (*Pseudomonas* I-F and I-E), and AcrIC1 (*Pseudomonas* I-C). In *Serratia*, CRISPR immunity provided an EOP of ~1×10^−6^ and the absence of EOP data indicates that no single plaques were detected with ~1×10^9^ pfu/mL of phage JS26. **(d)** Percentage distribution of the MGE origin (phage, plasmid/ICE, or unclear) for the collection of orthologs of each of the validated acrs. The isoelectric point (pI) and molecular weights (MW) of the validated acrs are shown.

Our screening revealed that each of the 9 candidate *acr* families tested inhibited the function of either one or more of the CRISPR-Cas subtype/variants challenged (Figure 2c). Most candidates potently inhibited the *Pectobacterium* and *Serratia* type I-F systems; only a few inhibited the *Serratia* type I-E and/or *Pseudomonas* type I-F, and none affected the function of the *Pseudomonas* type I-E and I-C systems. Overall, these results are consistent with the higher similarities between the type I-F and I-E CRISPR-Cas system variants present in the hosts encoding the *acrs* and the *Pectobacterium* and *Serratia* model system variants used for testing (Figure 2b). Interestingly, our results revealed that AcrIF18.1* and AcrIF22* exhibit broad inhibitory functions, robustly inhibiting the *Serratia* type I-E system and diverse type I-F variants (dual subtype inhibition denoted with an asterisk) (Figure 2b and 2c). Cross-subtype inhibitory activity has only been observed previously for AcrIIA5 (Hynes *et al.*, 2018; Marshall *et al.*, 2018), AcrVA3 (Marino *et al.*, 2018), and certain homologs of AcrIF6 (Pawluk, Staals, *et al.*, 2016). In addition to AcrIF18.1*, AcrIF15 strongly inhibited all three type I-F systems, whereas AcrIF16 and AcrIF17.1 inhibited *Pseudomonas* type I-F immunity less potently in our assays. AcrIE8 was the only type I-E-specific inhibitor identified in this work. All of the *acr* homologs tested (AcrIF20.2 - 65% identity, AcrIE8.2 - 76%, AcrIF18.2* - 96%, and AcrIF17.2 - 39%) showed comparable inhibitory activities to their counterparts, with the exception of AcrIF17.2 from *Citrobacter* which, unlike its distant homolog in *Pectobacterium*, did not inhibit the type I-F system from *P. aeruginosa* (Supplementary Fig S1).

The newly discovered Acr families present substantial differences in their biochemical properties. These include molecular weights (MW) ranging from 7.5 to ~20 kDa and varied predicted average isoelectric points (pI), spanning from acidic net charges (~pH = 4) to basic (~pH = 10) (Figure 2d, Supplementary Figure S2). Using a previously established CRISPRi system in PA14 I-F (Bondy-Denomy *et al.*, 2015; Pawluk *et al.*, 2017), we sought to investigate whether any of the newly identified *P. aeruginosa* I-F CRISPR-Cas system inhibitors (AcrIF15-18*) could impede host immunity at the target DNA binding stage. Our results indicate that the dual inhibitor AcrIF18* and AcrIF15 act as potent target DNA binding inhibitors of the surveillance complex (Supplementary Figure S3), while AcrIF16-17 did not block CRISPRi. The specific inhibitory mechanism of these proteins requires further investigation.

### Identified Acrs are spread across diverse Proteobacteria and MGE types

To explore the phylogenetic distribution of the new Acrs, publicly available prokaryotic sequences at NCBI were searched for homologs. The resulting analyses revealed a heterogeneous Acr distribution across Proteobacteria, most belonging to *Enterobacteriaceae* (Supplementary Figure S4, Supplementary Datasheet S4). As expected, the total collection of hosts encoding these *acrs* showed enrichment of genera frequently encoding CRISPR-Cas types I-F and/or I-E (e.g. *Salmonella*, *Serratia*, *Cronobacter*, *Klebsiella*, *Pectobacterium*) (Supplementary Datasheet S1). Interestingly, while AcrIF19-22 are primarily confined to species of the *Pectobacterium* genus, the rest (AcrIF15-IF18* and AcrIE8) radiate over wider phylogenetic ranges (Supplementary Figure S4, Supplementary Datasheet S4). For instance, homologs of AcrIF17 and AcrIF16 are present in bacterial families in addition to *Enterobacteriaceae*, such as *Pseudomonadaceae*, *Burkholderiaceae*, *Halomonadaceae*, and *Desulfobacteraceae*, and *Yersiniaceae*, *Vibrionaceae*, and *Shewanellaceae*, respectively.

We then scanned the genomic contexts (~25 kb upstream and downstream) surrounding the *acr* homologs to identify marker genes that could provide situational insights (Supplementary Datasheet S4). Our analyses revealed the association of the identified *acrs* with distinct types of MGEs, including phages and conjugative elements (Figure 2d). The position and composition of *acr* loci relative to neighboring gene cassettes, were highly variable between MGEs (Supplementary Figure S5). We observed related phage genomes harboring completely distinct *acr* loci, in some cases carrying a different *aca* gene (Figure 3, Supplementary Figure S5), and sometimes lacking a known *aca* (Supplementary Figure S6). For example, we found a number of sequenced *Pectobacterium* genomes with integrated prophages that have similarities with *Pectobacterium* phage ZF40 (a phage carrying the *acrIF8-aca2* locus) (Pawluk, Staals, *et al.*, 2016; Birkholz *et al.*, 2019), but in the remaining cases the prophage regions encoded different combinations of *acr*(s), together with *aca5* (Figure 3). This observation, together with the common detection of closely related *aca* and *acr* homologs within taxonomically mixed bacterial clades (Figure 1a, Supplementary Figure S4, Supplementary Datasheet S4), indicates that these genes are prone to frequent horizontal transfer between diverse MGEs.

**Figure 3.**
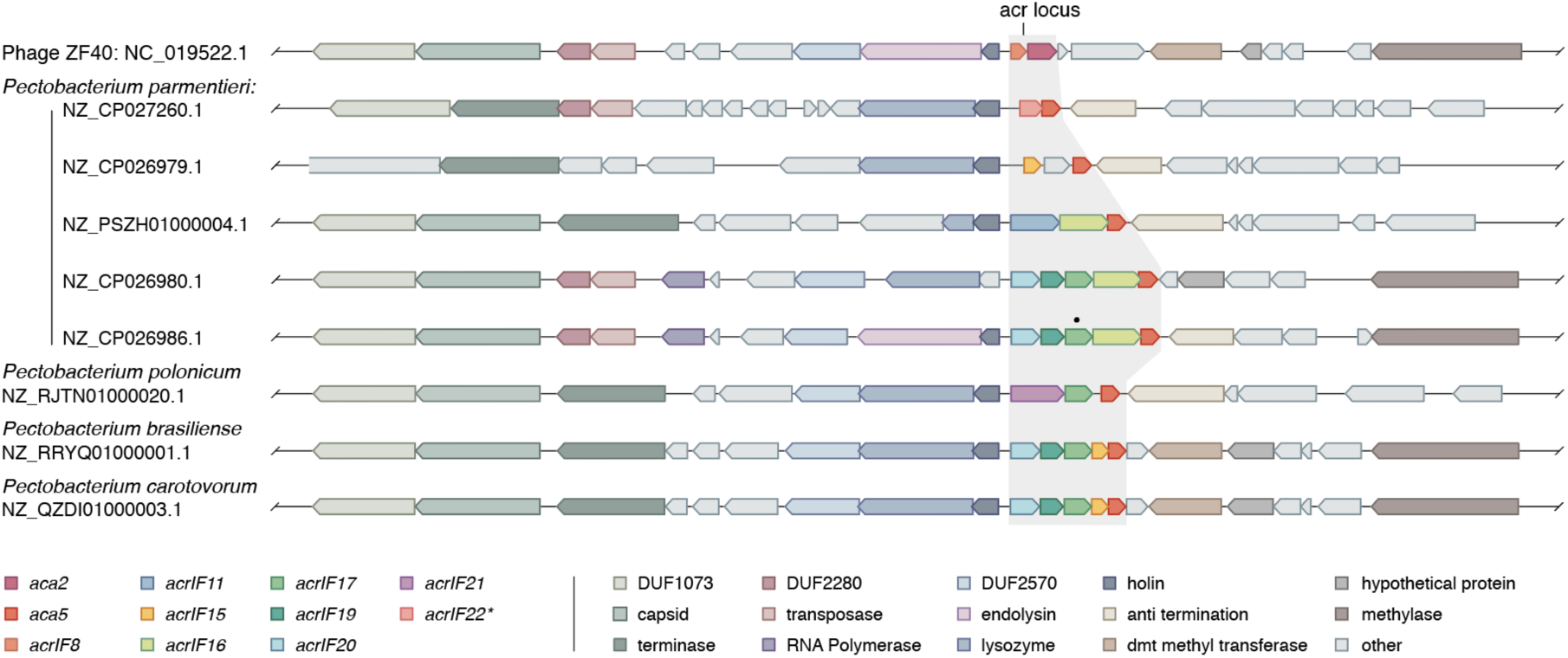
Genomic comparison between *Pectobacterium* phage ZF40 and different *Pectobacterium* genomes with related prophage/MGE regions. The alignment of the *acr* loci in related prophages/MGEs shows variability in nearby genes. Similar proteins are shown in the same color. Black dot denotes a pseudogene (early stop codon); domain of unknown function (DUF).

### Identification of a previously undescribed Aca and two type I-F Acrs

While exploring the genomic contexts of the identified *acrs*, we noticed that two AcrIF22* orthologs (encoded by a *Raoultella* phage and a *Klebsiella* plasmid) were located upstream of a small gene of hypothetical function distinct from, but in the same position as, *aca5* (Figure 4a). Motivated by this finding, we sought to investigate the potential Aca role of this gene further. A hallmark of all other previously identified Aca proteins is the presence of a helix-turn-helix (HTH) DNA-binding domain, necessary for the transcriptional repression of the *acr* operon (Birkholz *et al.*, 2019; Stanley *et al.*, 2019). Multiple sequence alignments of diverse representatives followed by functional domain prediction analyses revealed a conserved HTH motif in the hypothetical protein (Figure 4b, Supplementary Figure S7 and S8). Phylogenetic analyses showed a wide distribution of homologs across diverse Proteobacterial MGEs, including phages (50%) and plasmids (38%) (Figure 4c, Supplementary Datasheet S5). Moreover, our search revealed that this gene is also encoded downstream of an *acrIF15* homolog in a *Pectobacterium* plasmid (Figure 4c). Taken together, these results indicate this is an anti-CRISPR associated (*aca)* gene, hereafter referred to as *aca9*.

**Figure 4.**
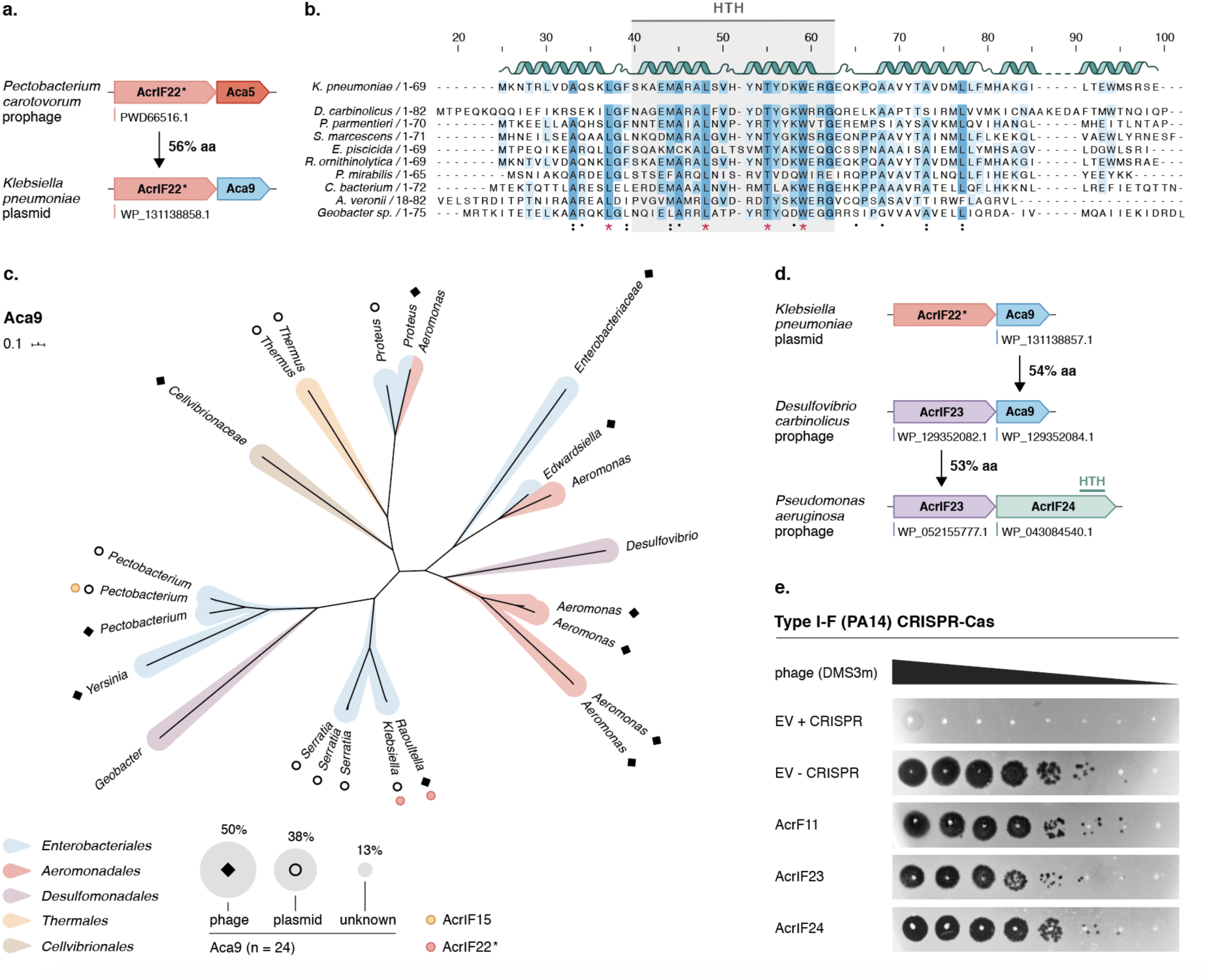
Discovery of Aca9 uncovers two potent I-F Acrs. **(a)** Identification of an AcrIF22* homolog reveals a previously undescribed *aca* marker gene (*aca9*). **(b)** Alignment of diverse Aca9 representatives carried by Proteobacterial MGEs. Predicted secondary structure alpha-helix regions are illustrated as ribbons for the *K. pneumoniae* homolog (top). The HTH DNA binding domain is highlighted in gray. **(c)** Phylogenetic diversity of Aca9 across publicly available sequences. Tree branches are colored by the bacterial host from where they originate (Order level). The MGE origin is shown with an empty circle (plasmid-like elements), a black diamond (phage), and left blank when the genomic context was unclear. The relative percentage distribution of *aca9* orthologs by MGE origin is depicted as the area of the gray shaded circles in the legend. The positions in the tree where *acrIF15* and *acrIF22\** are found associated with *aca9* are marked with yellow and red circles, respectively. **(d)** Guilt-by-association searches following *aca9* led to the discovery of two candidate *acr* genes in *P. aeruginosa*. **(e)** Functional testing for inhibition of the PA14 I-F CRISPR-Cas system was performed by phage plaque assays in strains carrying plasmids expressing the indicated *acr* candidates. Ten-fold serial dilutions of CRISPR-targeted DMS3m phage were titered on lawns of *P. aeruginosa* naturally expressing its I-F CRISPR-Cas system. The ΔCRISPR strain shows phage replication in the absence of CRISPR-Cas targeting. AcrIF11 was employed as a positive control for strong I-F inhibition.

To test whether *aca9* could be exploited as a marker for *acr* discovery, we searched for *aca9*-associated genes with homologs in *P. aeruginosa*, one of our model CRISPR-Cas organisms (Figure 4d). We identified a *P. aeruginosa* homolog of a hypothetical protein encoded next to *aca9* in *Desulfovibrio carbinolicus* that completely inactivated the PA14 type I-F CRISPR-Cas system (AcrIF23), as did its neighbor in a *P. aeruginosa* prophage (AcrIF24) (Figure 4e, Supplementary Figure S8). These two Acr proteins are homologous to a number of uncharacterized proteins that are distributed across Proteobacterial classes (Supplementary Figure S4). Although no known *aca* genes form part of the *P. aeruginosa acrIF23-24* locus, the C-terminus of AcrIF24 contains an HTH domain, implying a possible dual Acr-Aca function (Figure 4d, Supplementary Figure S8). A similar binary role was previously shown for AcrIIA1 (Osuna, Karambelkar, Mahendra, Sarbach, *et al.*, 2020) and AcrIIA13-15 (Watters *et al.*, 2020), suggesting that self-regulation by Acrs may be widespread. Our results reveal two additional I-F inhibitors associated with genes encoding proteins containing different HTH domains.

### Acrs cluster with Anti-RM and other anti-defense genes

A closer examination of the genomic environments surrounding the newly identified *acr* loci revealed several intriguing instances of additional anti-defense system components (Figure 5). For example, an anti-defense gene cluster was identified in a discrete region of a *Klebsiella pneumoniae* plasmid, separated from the gene modules responsible for plasmid housekeeping functions (e.g. replication, partitioning, and conjugative transfer) (Figure 5a). Together with an *aca9-acrIF22\** Acr locus, we found genes encoding anti-restriction modification systems (Anti-RM) (e.g. ArdA and KlcA (Zavilgelsky and Rastorguev, 2009; Serfiotis-Mitsa *et al.*, 2010)) and a plasmid SOS-inhibition (*psi*) locus involved in suppressing the deleterious host SOS response elicited by conjugative plasmid entry (Bagdasarian *et al.*, 1986). Orphan methyltransferase genes were also co-encoded in this region, suggesting a potential protective role against host restriction enzymes, as shown previously for other MGEs (Günthert and Reiners, 1987; Murphy *et al.*, 2013). Furthermore, *acrIF16* and *acrIF17* were found adjacent to a methyltransferase gene and in close proximity to an Anti-RM gene (*ardA*) in a *Rahnella* plasmid (Figure 5b(i)).

**Figure 5.**
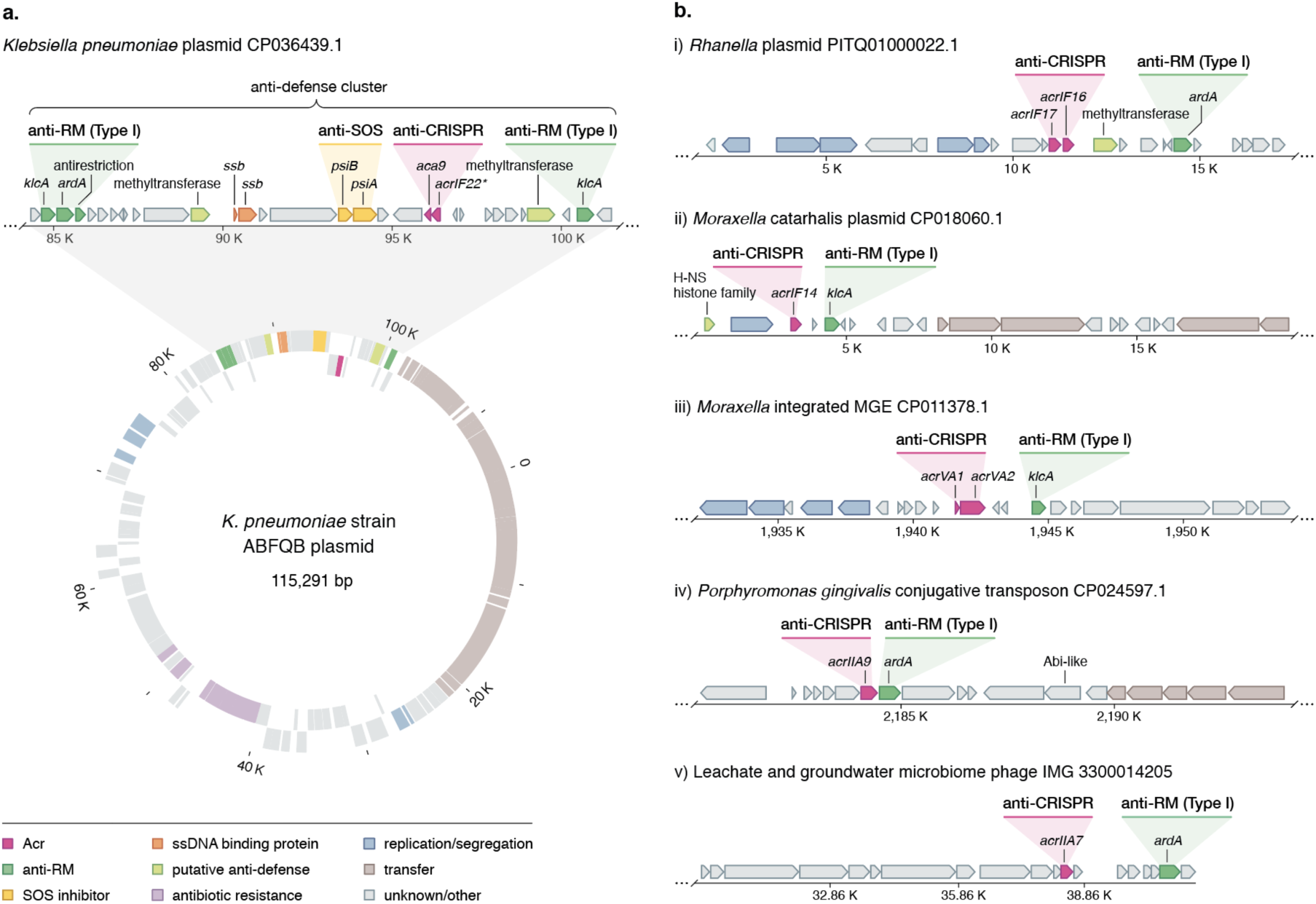
Acrs cluster with other antagonists of host defense functions; anti-defense islands. **(a)** Genome organization of the *Klebsiella pneumoniae* ABFQB plasmid unitig_1 (CP036439.1), highlighting the anti-defense cluster region. Genes are colored according to their predicted functions as shown in the key. Coding regions encoded on the plus and minus strands are shown on the outer and inner lanes of the gene map, respectively. **(b)** Examples of genomic regions within diverse MGEs where *acrs* were found in the vicinity of known anti-RM genes and other putative anti-defense determinants. Genes are colored according to the gene key in (a).

We then explored whether anti-RM genes are present nearby previously validated *acr* genes. Our analyses revealed the colocalization of the anti-RM gene *klcA* with previously described type I and V *acrs* (e.g. *acrIF14* and *acrVA1-2)* in *Moraxella* MGEs (Figure 5b(ii) and 5b(iii)). Moreover, a gene encoding an H-NS histone family homolog was adjacent to *acr* and anti-RM loci in a *Moraxella catarrhalis* plasmid (Figure 5b(iii)). MGE-encoded H-NS-like proteins are often considered “stealth proteins” that tamper with host gene expression (Shintani, Suzuki-Minakuchi and Nojiri, 2015; Vial and Hommais, 2019). Given the H-NS-mediated silencing of CRISPR-Cas adaptive immunity in *E. coli* (Pul *et al.*, 2010; Westra *et al.*, 2010), the lateral acquisition of H-NS homologs may help MGEs evade adaptive immunity, as previously proposed for certain plasmids (Dorman, 2014; Dorman and Ní Bhriain, 2020) and phages (Skennerton *et al.*, 2011). Finally, we also found close ties between type II *acrs* (e.g. AcrIIA9 and AcrIIA7) and anti-RM components in other types of MGEs, including a conjugative transposon and a putative phage (Figure 5b(iv) and 5b(v)).

Collectively, these results suggest that, apart from accumulating *acrs* against CRISPR-Cas systems (Figure 1 to 4), MGEs compile a broader arsenal of inhibitors to overcome other host immune mechanisms. Intriguingly, the resulting collection of inhibitors appears to mirror the clustered arrangement of defense systems in their hosts, termed “defense islands” (Makarova *et al.*, 2011; Doron *et al.*, 2018). We found that the gene neighborhoods of loci encoding Acr and Anti-RM proteins are typically crowded with other small hypothetical protein-coding genes of unknown function (Figure 5). We speculate that these “anti-defense islands” may constitute an unrecognized phenomenon in diverse MGEs, potentially enriched with new genes that antagonize diverse defense systems (Bernheim and Sorek, 2019).

## Discussion

Following an *aca*-based guilt-by-association search, 11 Acr families and a new Aca family were discovered across chromosomal and extrachromosomal MGEs of mostly Enterobacteriaceae (Figure 1b and 4d). These findings ascribe function to a dozen gene families that were previously only hypothetical and reveals that the diverse MGEs that carry them are likely encountering and evading functional CRISPR immunity *in situ.* The Acr proteins identified share no sequence homology with known Acrs, increasing the collection of distinct subtype I-F Acrs to 24 (from 14) and I-E CRISPR-Cas inhibitors to 10 (from 7). Many genomes analyzed showed instances of self-targeting, where integrated MGEs carrying the *acr* operons had regions with perfect identity to host-derived CRISPR spacers (Figure 1b and c). One self-targeting spacer was even predicted to target an *acr* gene *(acrIF16*), encoded by a *Pectobacterium parmentieri* prophage (Figure 1c, bottom, Supplementary Datasheet S2).

Loci encoding Acrs frequently contain more than one *acr* gene (Bondy-Denomy *et al.*, 2013; Pawluk *et al.*, 2014; Rauch *et al.*, 2017; Marino *et al.*, 2018). Here we observed loci where as many as 4 distinct *acrs* are “stacked” upstream of *aca5* (Figure 1). It is unclear what fitness benefits are associated with such locus organizations and whether functional redundancy or cooperation between the different Acrs occurs. Given the fast MGE-host co-evolutionary arms race, carrying multiple *acrs* likely serves as a safeguard against Cas mutational escape or subtype diversity. Alternatively, multiple Acr proteins could provide MGEs with “division of labor” potential where different Acr proteins are used during distinct stages of the MGE life cycle, or contribute to a more robust inhibitory effect by blocking the immune pathway at different stages (Osuna, Karambelkar, Mahendra, Christie, *et al.*, 2020).

Former mechanistic characterizations of Acr proteins have revealed a remarkable diversity of inhibitory functions (Davidson *et al.*, 2020). Interestingly, relatively low MWs and pIs have been reported for Acrs inhibiting CRISPR-Cas systems via DNA mimicry (Jiang *et al.*, 2019; Liu *et al.*, 2019), suggesting a putative link between low MW/pI values and mechanistic inhibitory function. Given the small size and negative charges of AcrIF18* and AcrIF15 (Supplementary Figure S2), and their Cascade DNA binding inhibitory activities (Supplementary Figure S3), it is possible that they function as DNA mimics – analogous to AcrIF2 (Chowdhury *et al.*, 2017), AcrIF10 (Guo *et al.*, 2017), AcrIIA2 (Shin *et al.*, 2017; Liu *et al.*, 2019) and AcrIIA4 (Dong *et al.*, 2017). DNA-recognition domains in Cas proteins (and components of other bacterial defense systems) are evolutionarily constrained, making them desirable targets for broad spectrum inhibition. A great example of how MGEs exploit this “Achilles heel” is phage protein Ocr, a DNA mimic that provides protection from both type I RM systems and BREX (Bacteriophage exclusion systems) (Isaev *et al.*, 2020). While DNA mimicry could help explain the broad inhibitory activity of AcrIF18*, further experiments are required to ascertain such functionality.

Although the Acr families we describe here are predominantly detected on prophage elements (Figure 2d, 3 and 4c), many are also carried by other types of MGEs, such as conjugative plasmids and ICEs (Figure 2d and 4c). Our results support the notion that Acrs play an important role in facilitating the horizontal transfer of diverse MGE-encoded traits, such as plasmid-encoded antibiotic resistance determinants (Figure 5, Supplementary Datasheet S6) (Mahendra *et al.*, 2020). Consistent with this idea, we recently reported a positive association of *acr* genes and acquired antibiotic resistance genes in *P. aeruginosa* genomes (Shehreen *et al.*, 2019). Interestingly, the taxonomically broadly distributed AcrIF17 is particularly enriched on conjugative plasmids (Figure 2d). Because conjugative elements often exhibit broader transfer host ranges than phages (Norman, Hansen and Sørensen, 2009; Guglielmini *et al.*, 2011) these results may reveal a relationship between the MGE origin of *acrs*, their phylogenetic distribution and inhibitory spectrum.

Consistent with previous work, we show that Acrs tend to inhibit the specific CRISPR-Cas system(s) present in the hosts of the MGEs carrying them, although the inhibition spectrum of certain Acrs is occasionally broader (Pawluk, Staals, *et al.*, 2016; Marino *et al.*, 2018; Forsberg *et al.*, 2019; Song *et al.*, 2019). These data indicate the challenge of inferring inhibitory activity *a priori* and highlights the necessity to interrogate *acr* function experimentally in a case by case manner using a panel of CRISPR-Cas systems.

Because the dynamics of gene flow within microbial communities are governed by the interactions between MGEs and their hosts, shedding light on the defense/anti-defense arms race is integral for understanding the ecology and evolution of bacteria. However, prokaryotes possess an extraordinary variety of defense mechanisms and identifying uncharted immune systems has proved challenging. Previous work has shown that defense systems often co-localize within defense islands in bacterial genomes (Makarova *et al.*, 2011), thus allowing the identification of undiscovered immune systems (Doron *et al.*, 2018; Cohen *et al.*, 2019). Here, we find that *acr* loci often cluster in MGE genomes with antagonists of other host defense functions (e.g. anti-RM), indicating that MGEs organize their counter defense strategies in “anti-defense islands”. We anticipate that this genetic co-occurrence will be useful for the discovery of novel anti-defense systems.

## Methods

### Bioinformatic searches and phylogenetic analyses of Aca and Acr proteins

Protein sequences of Aca5 homologs were identified through 4 iterations of PSI-BLAST with default search parameters against the non-redundant protein database (NCBI-NR) using the *Pectobacterium* Aca5 WP_039494319.1 sequence as a query. Only hits with >70% coverage and e-value <10^−8^ were included in the generation of the position-specific scoring matrix (PSSM). An HMM model of Aca5 was also built using PSI-BLAST hits (three iterations; query cover: >90% and identity >55%). Acr candidates were identified following a previously described guilt-by-association approach (Pawluk, Staals, *et al.*, 2016). Briefly, hypothetical ORFs upstream genes encoding Aca5 homologs were found through a combination of bioinformatic searches using PSI-BLAST (up to three iterations, only considering hits with e-values <10^−4^ for PSSM generation) and hmmsearch (HMMER v3.0; e-value cutoff 0.05; the script was modified to extract upstream and downstream nucleotide sequences for each hit).

Multiple alignments of the identified Aca and Acr proteins were performed with the MUSCLE software (Edgar, 2004). Alignment sites with gaps were trimmed from the alignment. Maximum Likelihood phylogenetic trees were constructed using MEGA (Kumar, Stecher and Tamura, 2016) (500 bootstraps and standard settings) and displayed using iTOL (Letunic and Bork, 2019).

### MGE/prophage context, CRISPR-Cas and self-targeting analyses

The genomic contexts (<25 Kb upstream and downstream) of all identified *acr* homologs were scanned manually in search for prophage, plasmid, and ICE signature protein-coding genes. The MGE type and corresponding signature gene(s) used to determine the *acr* origin are displayed in Supplementary Datasheets S1. In the absence of annotated genes, PSI-BLAST searches (70% coverage, < 10^−4^ e-value) were performed with genes neighboring the *acrs* in an attempt to find homology to genes of known function that could provide situational insights. CRISPRCasFinder (Couvin *et al.*, 2018) was employed to determine the presence and sequence integrity of the CRISPR-Cas systems in the genomes of the bacterial hosts encoding the selected *acr* candidates and to extract the corresponding CRISPR spacers (Supplementary Datasheets S1 and S4). Self-targeting analyses were performed using CRISPRTarget (Biswas *et al.*, 2013), STSS (Watters *et al.*, 2018), and CRISPRminer(v2) (Zhang *et al.*, 2018) by blasting the spacer contents against the host genome assemblies (Supplementary Datasheet S2). Prophage regions were identified and annotated via PHASTER (Arndt *et al.*, 2016) and the position of the *acr* loci and self-targets were compared to the predicted prophage regions (Supplementary Datasheet S2).

### CRISPR-Cas sequence identity comparisons

The protein sequences of the Cas orthologs of the model CRISPR-Cas systems tested and the endogenous CRISPR-Cas systems (hosts from which the selected *acrs* originate) were aligned using BLASTp. An average score of the percentage sequence identity between systems was calculated (Supplementary Datasheet S3). Only the proteins involved in the interference complex were taken into consideration in this analysis (i.e. excluding the adaptation module: Cas1 and Cas2 in I-E systems and Cas1 in I-F systems).

### Bacterial strains, phages and growth conditions

All strains and phages used in this study are listed in Supplementary Tables S1 and S2. *Pseudomonas aeruginosa* strains (PA14, PAO1) and *Escherichia coli* strains (Mach-1) were routinely grown at 37°C in Lysogeny Broth (LB) (10 g/L tryptone, 5 g/L yeast extract, and 10 g/L NaCl). *Pectobacterium atrosepticum* SCRI1043 and *Serratia* sp. ATCC39006 were grown at 25°C and 30°C, respectively in LB (10 g/L tryptone, 5 g/L yeast extract, and 5 g/L NaCl). All solid plate media were supplemented with 1.5% w/v agar. Media were supplemented with antibiotics to maintain the pHERD30T plasmid (and derivatives): 30 µg/mL for *Pectobacterium atrosepticum* SCRI1043 and *Serratia* sp. ATCC39006, 15 µg/mL gentamicin for *E. coli*, and 50 µg/mL gentamicin for *P. aeruginosa*. When appropriate, the following inducer concentrations were used: 0.1-0.3% w/v arabinose and 0.1 mM isopropyl β-D-1-thiogalactopyranoside (IPTG). During heat shock transformations, *E. coli* was recovered in SOC media (20 g tryptone, 5 g yeast extract, 10 mM NaCl, 2.5 mM KCl, 10 mM MgCl_2_, 10 mM, MgSO_4_, and 20 mM glucose in 1 L dH_2_O).

*Pseudomonas* phages DMS3, DMS3m, and JBD30 derivatives were propagated on PA14 ΔCRISPR or PAO1 WT. *Pectobacterium* phage ϕTE (Blower *et al.*, 2012) and *Serratia* phage JS26 (Jackson *et al.*, 2019) were propagated on *P. atrosepticum* SCRI1043 (WT) (Bell *et al.*, 2004) and *Serratia* sp. ATCC39006 LacA strains (Thomson *et al.*, 2000), respectively. *Pseudomonas* phages were stored at 4°C in SM buffer (100 mM NaCl, 8 mM Mg2SO_4_, 50 mM Tris-HCl pH 7.5, 0.01% w/v gelatin) over chloroform. *Pectobacterium* and *Serratia* phages were stored at 4°C in phage buffer (10 mM Tris-HCl pH 7.4, 10 mM MgSO_4_ and 0.01% w/v gelatin) over chloroform.

### Cloning of candidate anti-CRISPR genes

Candidate *acr* genes identified in the *aca5*-based guilt-by-association search (Supplementary Datasheet S1) were synthesized as gene fragments (Twist Biosciences) and cloned into the NcoI and HindIII sites of the pHERD30T shuttle vector using Gibson Assembly (New England Biolabs). The plasmid constructs were propagated in commercial *E. coli* Mach-1 competent cells (Invitrogen, Thermo Fisher Scientific) upon transformation, following the manufacturers recommendations. AcrIE8.1, AcrIF17.2, and AcrIF18.2* were synthesized as gBlocks (IDT) and ligated into the NcoI and HindIII sites of the pHERD30T shuttle vector and transformed into *E. coli* DH5α competent cells. The integrity of the cloned fragments were verified via Sanger sequencing using primers outside of the multiple cloning site (Supplementary Table S3). A list of the plasmids and oligonucleotides used in this study can be found in Supplementary Table S3 and S4.

### Preparation of *P. aeruginosa*, *P. atrosepticum*, and *Serratia* sp. ATCC39006 strains for *acr* functional testing

The pHERD30T plasmids with different candidate anti-CRISPRs were electroporated into the different *P. a*eruginosa strains. Briefly, overnight cultures were washed twice in 300 mM sucrose and concentrated tenfold. Competent cells were then transformed with 50 – 100 ng plasmid and incubated without plasmid selection in LB broth for 1 hr at 37 °C before they were grown overnight at 37°C on LB agar plates with plasmid selection. For the transformation of these plasmids in *P. atrosepticum* and *Serratia* ATCC39006, they were first transformed into the chemical competent *E. coli* ST18 and cells were plated onto LB agar plates with 50 µg/mL 5-aminolevulinic acid (ALA). Then, the donor ST18 cells were conjugated with recipient *P. atrosepticum* (PCF188) (Pawluk et al. 2016) and *Serratia* ATCC39006 PCF524 (I-E) and PCF525 (I-F). The positive clones were selected by streaking onto LB agar plates containing 30 µg/mL gentamicin. The arabinose-inducible promoter in pHERD30T was used to drive the expression of the candidate *acr* genes.

### *P. aeruginosa* phage immunity assays

The functionality of the identified *acr* candidate genes was assessed through phage spotting assays or efficiency of plaquing (EOP). These tests evaluated the replication of CRISPR-targeted phages DMS3m (I-F) and JBD30 (I-C and I-E) in bacterial lawns relative to the empty vector control. Efficiency of plaquing (EOP) was calculated by dividing the number of plaque forming units (pfus) formed on a phage-targeting strain by the number of pfus formed on a non-targeting strain: ΔCRISPR strain (I-F and I-E) or the absence of CRISPR expression inducer (I-C). Additional controls included infection in the presence of an already validated Acr (i.e. AcrIE4/F7, AcrIF11, or AcrIC1). Each pfu calculation was performed in 3 biological replicates and expressed as the mean EOP +/− SD (error bars). Briefly, 200 µL of bacterial overnight cultures were mixed with 10 µL of ten-fold phage serial dilutions and combined with 4 mL of molten top agar (0.7%) supplemented with 10 mM MgSO_4_. The mix was poured onto LB agar (1.5%) plates containing 50 µg/mL gentamicin and 10 mM MgSO_4_. Additionally, plates were supplemented with 1 mM IPTG and 0.3% w/v arabinose for experiments using the PAO1 strain (I-C CRISPR-Cas) and with 0.3% w/v arabinose when using the PA4386 strain (I-E CRISPR-Cas), and PA14 strain (I-F CRISPR-Cas). Plates were incubated overnight at 30°C and phage plaque-forming units (pfu) were calculated. In phage spotting experiments, phage dilutions 3.5 µL of ten-fold serial dilutions of the phage lysates were spotted onto the plate surface containing the bacterial lawn in the top agar. Plate images were obtained using Gel Doc EZ Gel Documentation System (BioRad) and Image Lab (BioRad) software.

### Anti-CRISPR Assay in *P. atrosepticum* and *Serratia* sp. ATCC39006

PCF188, PCF524, and PCF525 were transformed with plasmids expressing the different Acrs and overnight cultures were used to pour top agar plates (100 µL culture added to 4 mL LB with 0.35% agar) onto LB agar plates (containing 30 µg/mL gentamicin and 0.1% w/v arabinose). Due to the toxic effects on *Serratia* cells, AcrIF9 (positive control for inhibition) was tested in the absence of arabinose. Twelve-fold serial dilutions of phage ϕTE and JS26 lysates were made in phage buffer, and 15 µL was spotted on the dried top agar plates. Then, the plates were incubated overnight at 25°C for *Pectobacterium* and 30°C for *Serratia*. The efficiency of plaquing (EOP) was determined by calculating the pfus per mL of the *Pectobacterium* 3xTE and *Serratia* cells (expressing the different Acrs) divided by the pfus of the corresponding wild-type with an empty vector pHERD30T control. Three biological replicates were done for each of the experiments (Supplementary Datasheet S9).

### Construction of recombinant *acr* phages

DMS3m phage derivatives encoding AcrIF15, AcrIF18*, AcrIF16, and AcrIF17 were constructed. Briefly, a recombination plasmid with homology to the DMS3m acr locus (Borges *et al.*, 2018) was used to clone the Acr genes of interest upstream of *aca1* by Gibson Assembly (New England Biolabs). The resulting vectors were used to transform PA14 ΔCRISPR and the strains were infected with WT DMS3m::gent35 cassette. Phages were recovered after full plate infections in selection plates containing 50 µg/mL gentamicin and the resultant phage lysates were used for full plate infections in plates without selection. Recombinant phages were sequentially passaged through PA14 ΔCRISPR 3 times to purge away potential non-recombinant phage carryover. The integrity of the cloned Acr genes were verified by Sanger sequencing. Phages were stored in SM buffer at 4°C.

### Generation of recombinant Acr phage PA14 (dCas3) lysogens

200 µL of PA14dCas3 overnight culture was added to 4 mL of 0.7% LB top agar and spread on 1.5% LB agar plates supplemented with 10 mM MgSO_4_. 3 µL of the recombinant *acr* phages were spotted on the top agar bacterial lawns and plates were incubated at 30°C overnight. Following incubation, bacterial survivors (lysogens which undergo superinfection exclusion or are surface receptor mutants) within the plaques were isolated and spread on 1.5% LB agar plates. Single colonies were assayed for phage resistance by streaking across a line of phage lysate, compared to a sensitive WT PA14 control. Prophages were confirmed by sequencing, checking for resistance to superinfection by the same phage, and assessing the spontaneous production of phage from the lysogenic strain (supernatant was titered on PA14 ΔCRISPR).

### CRISPRi-based pyocyanin repression assay in *P. aeruginosa*

The pyocyanin repression assay was carried out as described previously (Bondy-Denomy *et al.*, 2015). Briefly, the PA14dCas3 lysogens harboring the constructed DMS3m prophages expressing either AcrIF15, AcrIF16, AcrIF16 or AcrIF18* were transformed with pHERD30T::crRNA*phzM*, a vector expressing a crRNA designed to target the promoter of *phzM*, a chromosomal gene in PA14 which is involved in the production of the pyocyanin (blue-green pigment). In the presence of crRNA *phzM* CRISPRi repression leads to a color change in the culture medium, from green to yellow. The different lysogens were additionally transformed with an empty vector (pHERD30T), serving as a non-targeting control (remains green).

Cultures of 3 independent lysogens were grown overnight in LB supplemented with gentamicin (50 µg/mL) for vector retention and 0.3% arabinose to induce crRNA expression. Pyocyanin was extracted from the overnight cultures and quantified by measuring absorbance at 520 nm, as described previously (Bondy-Denomy et al., 2015). Representative pictures of the color changes are also displayed (Supplementary Figure S10).

### Software and statistical analysis

Numerical data were analyzed and plotted with GraphPad Prism 6.0 Software. Statistical parameters are specified in the figure legends. HHPred searches were carried out for the prediction of protein domains (e.g. HTH). Protein secondary structure predictions were carried out by JPred4 (Drozdetskiy *et al.*, 2015) and Phyre2 (Kelley *et al.*, 2015) of antibiotic resistance genes was performed via BLAST against the Comprehensive Antibiotic Resistance Database (CARD) (McArthur *et al.*, 2013) (Supplementary Datasheet S6). A list of the Software is provided in Supplementary Table S5.

## Supplementary information

**Table S1.**
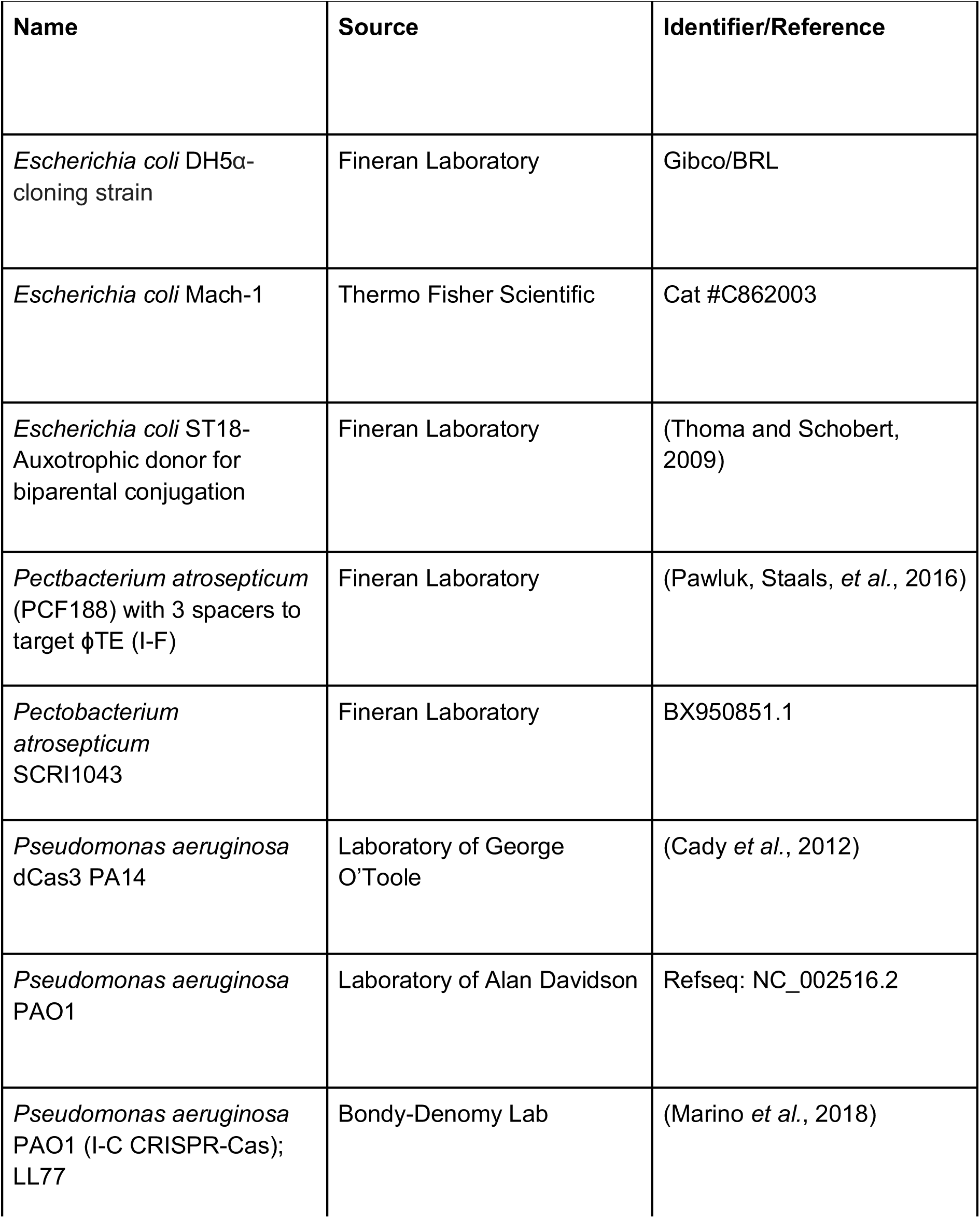
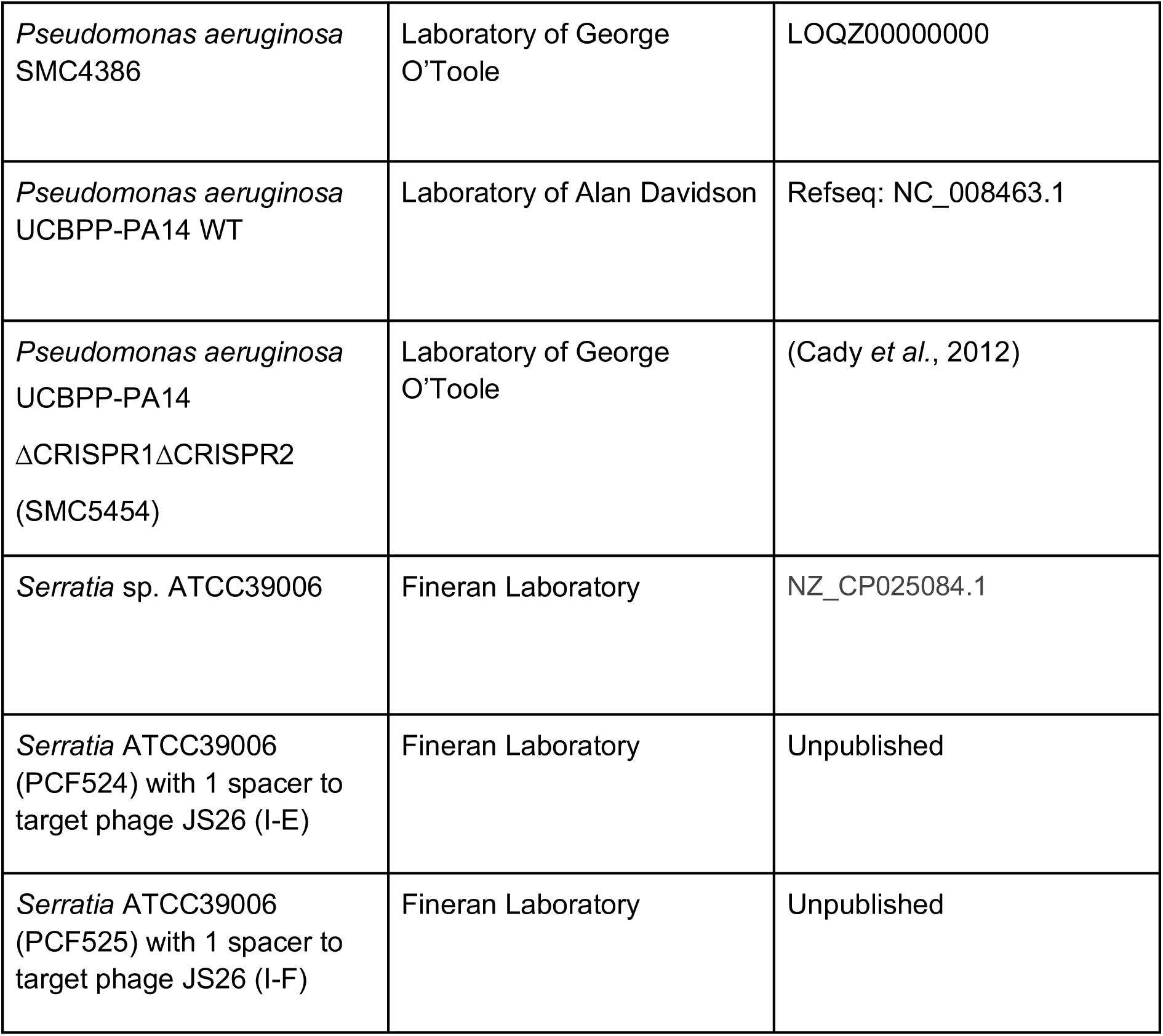
Bacterial strains used in this study

| Name | Source | Identifier/Reference |
| --- | --- | --- |
| *Escherichia coli* DH5α-cloning strain | Fineran Laboratory | Gibco/BRL |
| *Escherichia coli* Mach-1 | Thermo Fisher Scientific | Cat #C862003 |
| *Escherichia coli* ST18-Auxotrophic donor for biparental conjugation | Fineran Laboratory | (Thoma and Schobert, 2009) |
| *Pectobacterium atrosepticum* (PCF188) with 3 spacers to target ϕTE (I-F) | Fineran Laboratory | (Pawluk, Staals, *et al.*, 2016) |
| *Pectobacterium atrosepticum* SCRI1043 | Fineran Laboratory | BX950851.1 |
| *Pseudomonas aeruginosa* dCas3 PA14 | Laboratory of George O'Toole | (Cady *et al.*, 2012) |
| *Pseudomonas aeruginosa* PAO1 | Laboratory of Alan Davidson | Refseq: NC_002516.2 |
| *Pseudomonas aeruginosa* PAO1 (I-C CRISPR-Cas); LL77 | Bondy-Denomy Lab | (Marino *et al.*, 2018) |
| *Pseudomonas aeruginosa*<br>SMC4386 | Laboratory of George<br>O'Toole | LOQZ00000000 |
| *Pseudomonas aeruginosa*<br>UCBPP-PA14 WT | Laboratory of Alan Davidson | Refseq: NC_008463.1 |
| *Pseudomonas aeruginosa*<br>UCBPP-PA14<br>ΔCRISPR1ΔCRISPR2<br>(SMC5454) | Laboratory of George<br>O'Toole | (Cady *et al.*, 2012) |
| *Serratia* sp. ATCC39006 | Fineran Laboratory | NZ_CP025084.1 |
| *Serratia* ATCC39006<br>(PCF524) with 1 spacer to<br>target phage JS26 (I-E) | Fineran Laboratory | Unpublished |
| *Serratia* ATCC39006<br>(PCF525) with 1 spacer to<br>target phage JS26 (I-F) | Fineran Laboratory | Unpublished |

**Table S2.** Bacteriophages used in this study

| Name | Source | Identifier |
| --- | --- | --- |
| *Pectobacterium* Phage ϕTE | Fineran Laboratory | (Blower *et al.*, 2012) |
| *Pseudomonas* phage JBD30 | Laboratory of Alan Davidson | Refseq: NC_020298.1 |
| *Pseudomonas* phage<br>DMS3m (DMS3 derivative) | Laboratory of George<br>O'Toole | Refseq (DMS3):<br>NC_008717.1 |
| *Pseudomonas* phage<br>DMS3m<sub>acrIF15-18*</sub> | Bondy Denomy Laboratory | This study |
| *Serratia* Phage JS26 | Fineran Laboratory | (Jackson *et al.*, 2019) |

**Table S3.**
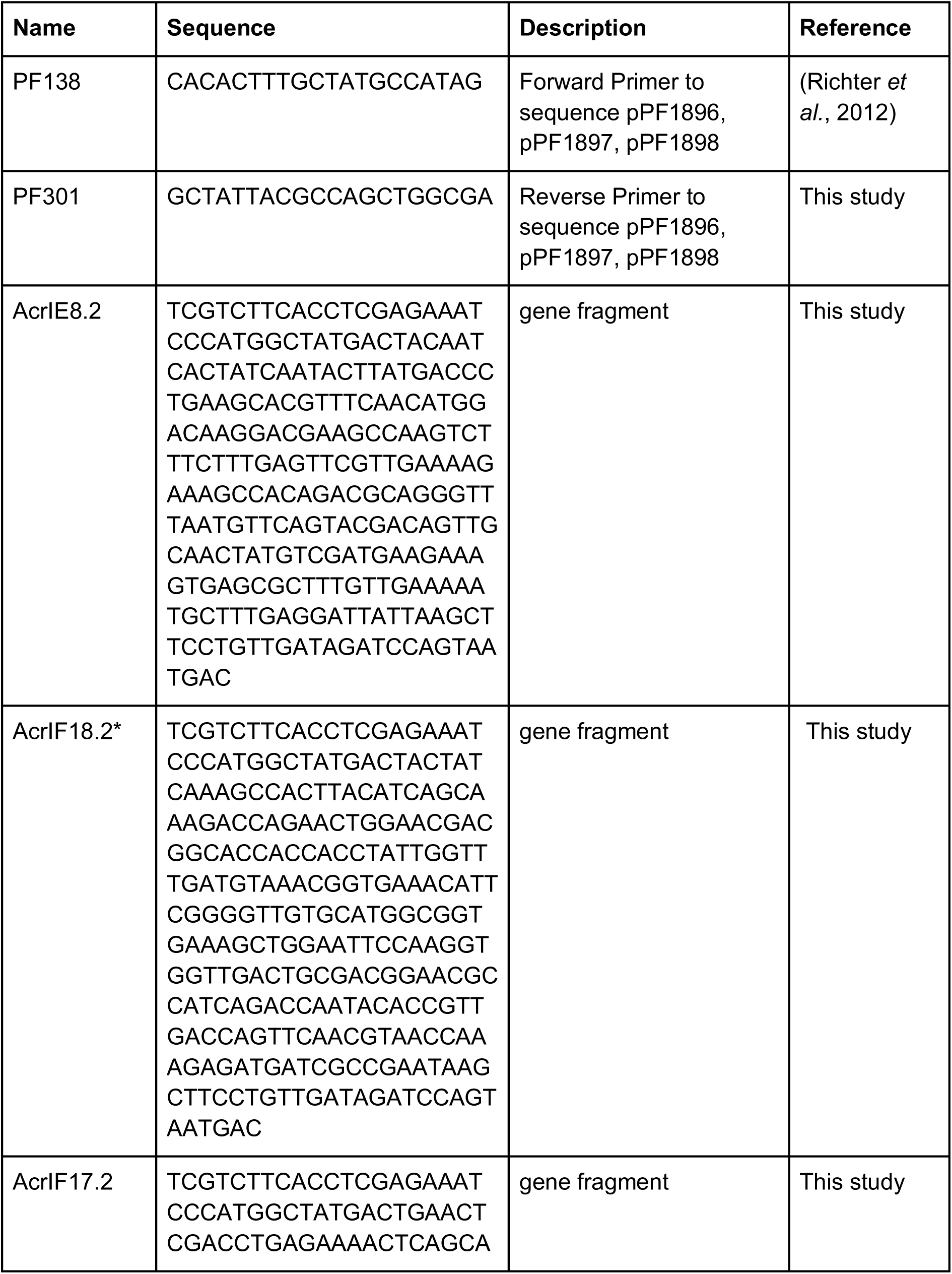
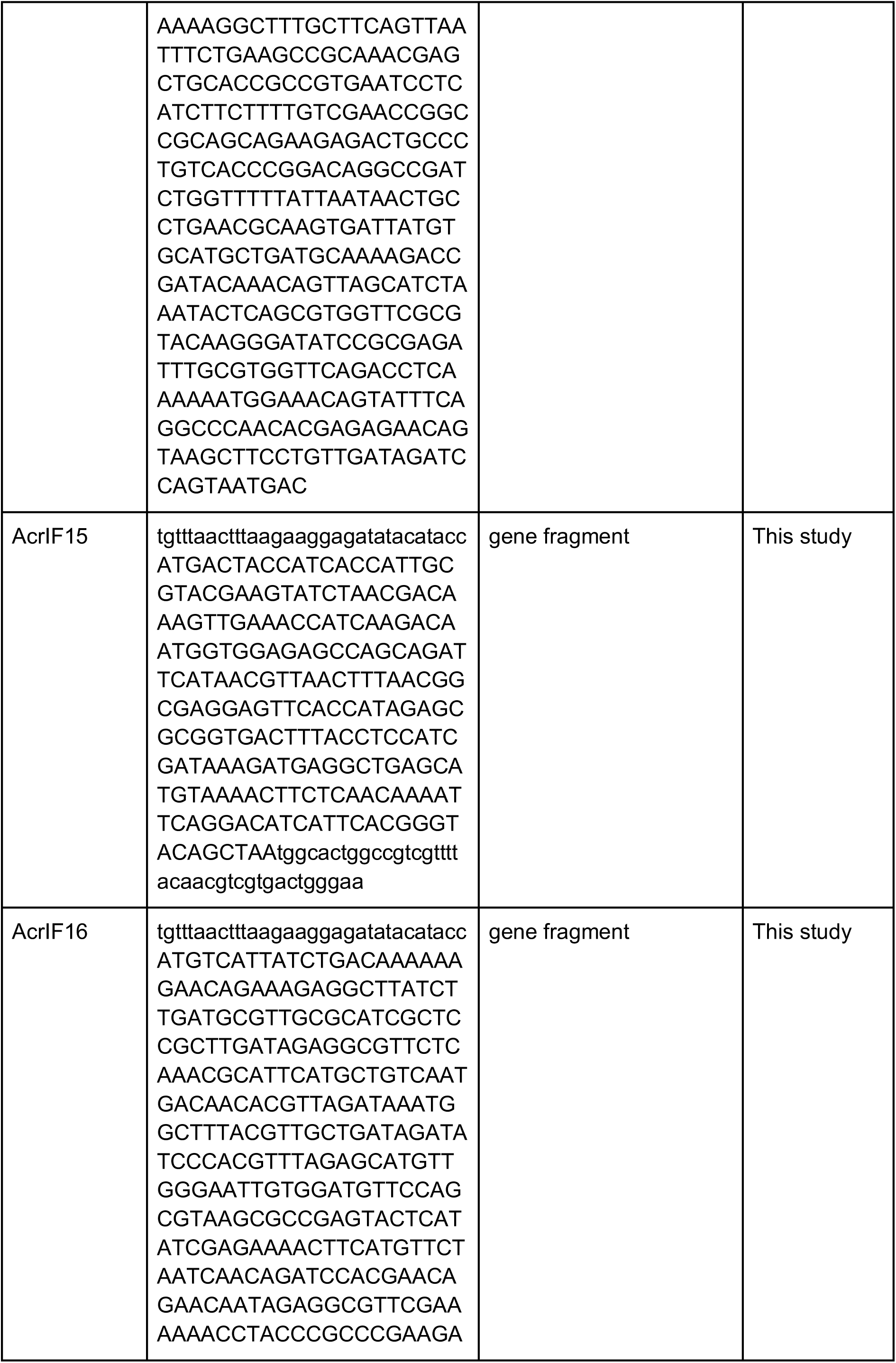
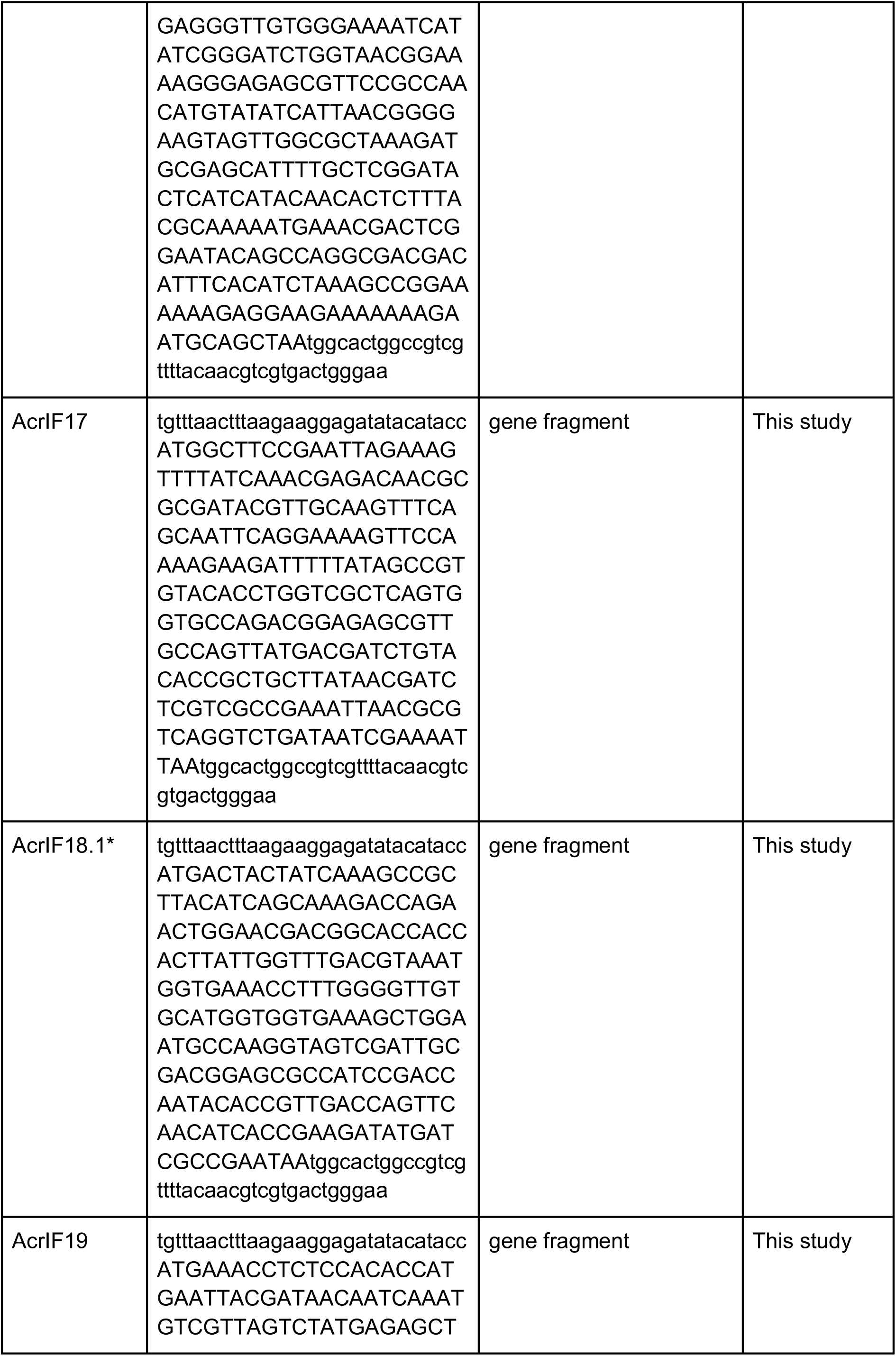
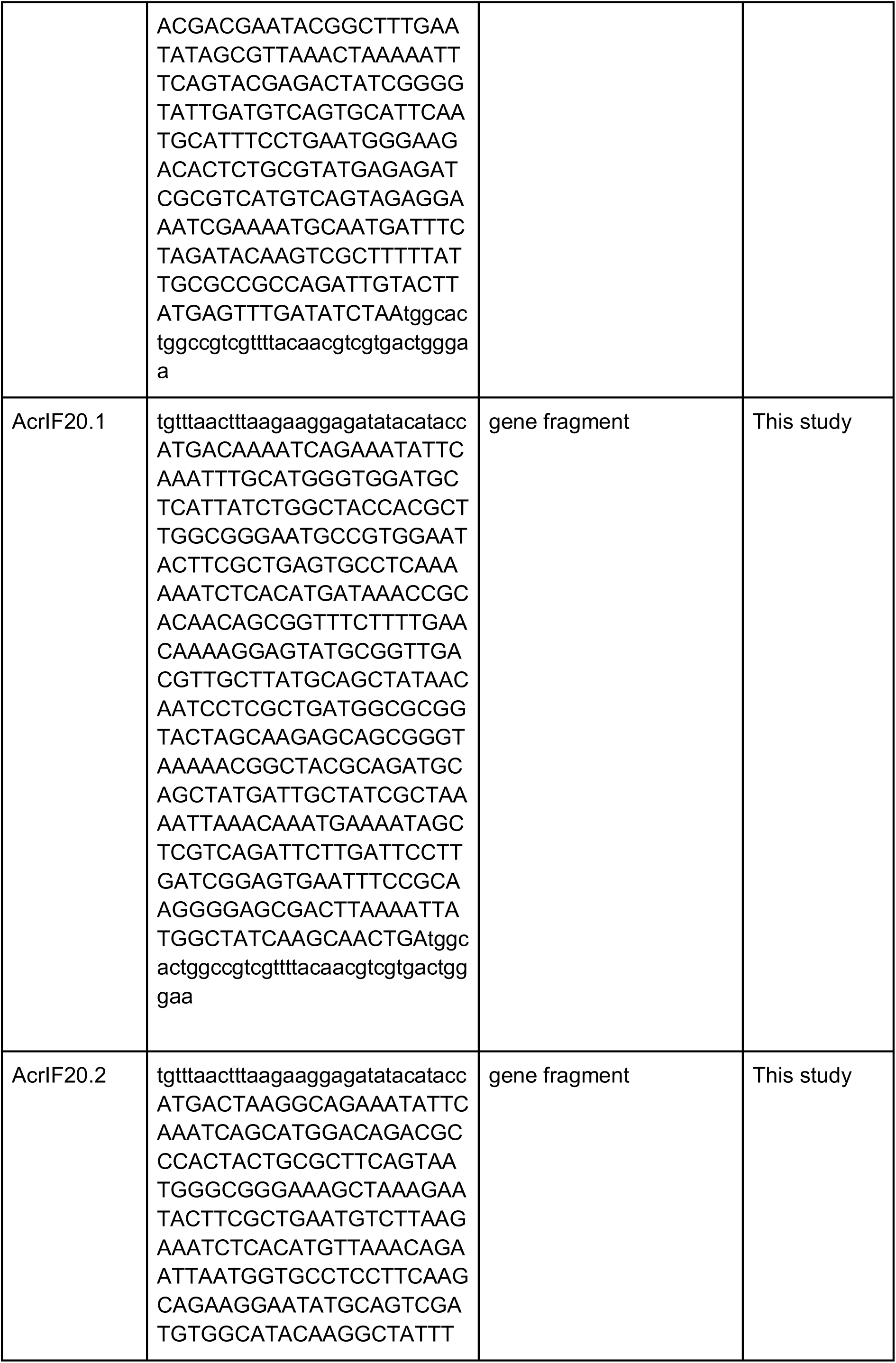
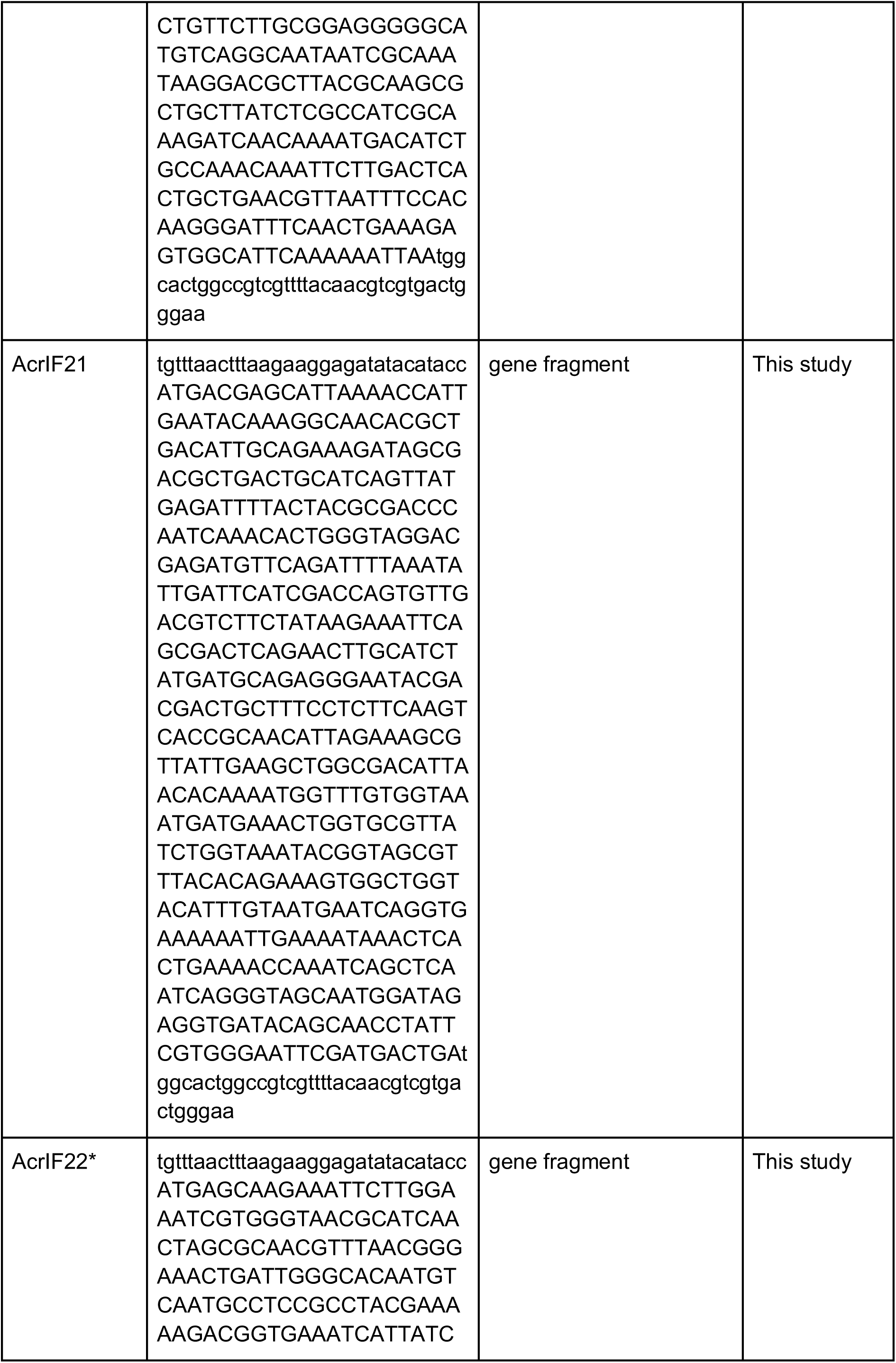
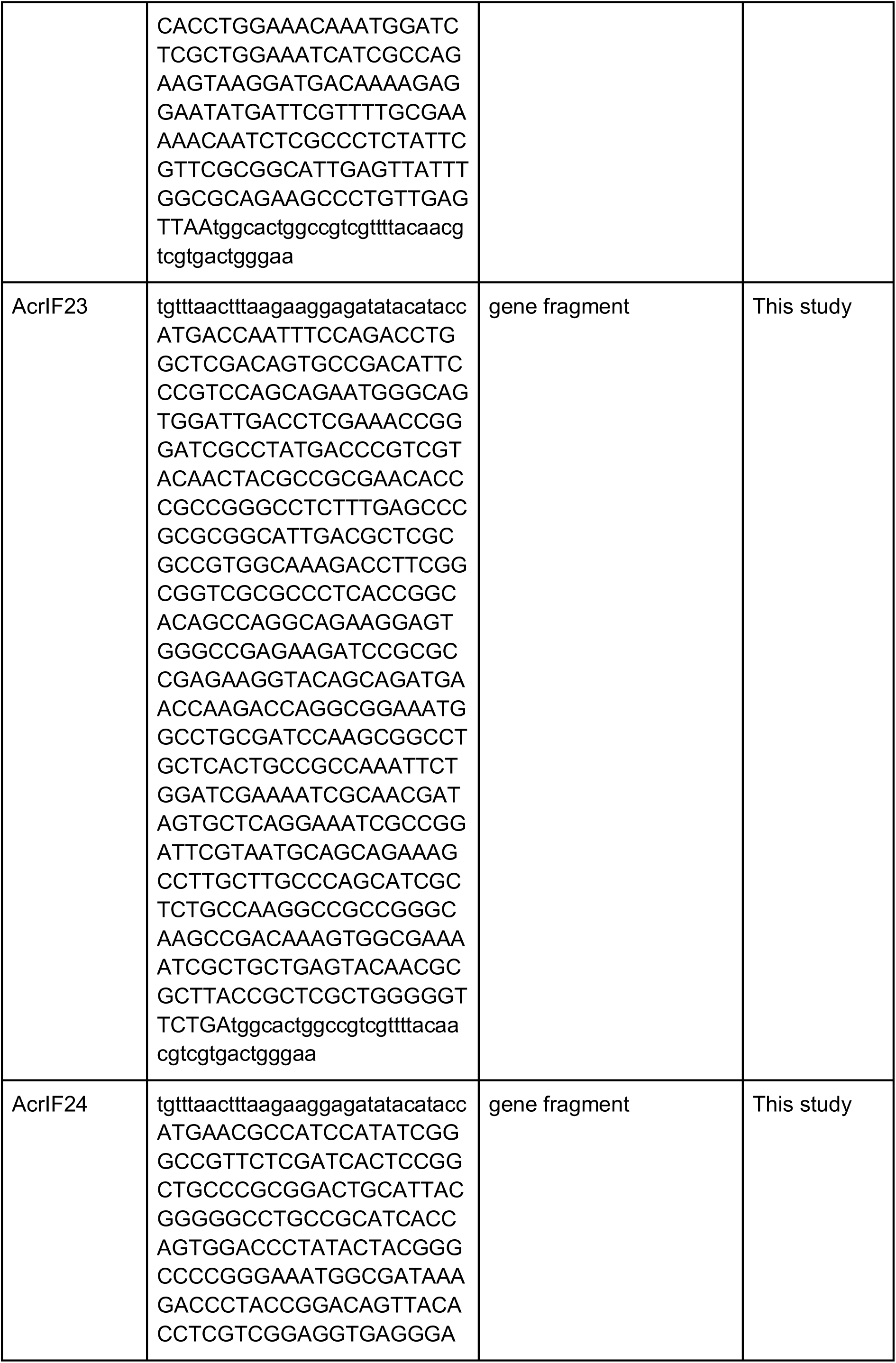
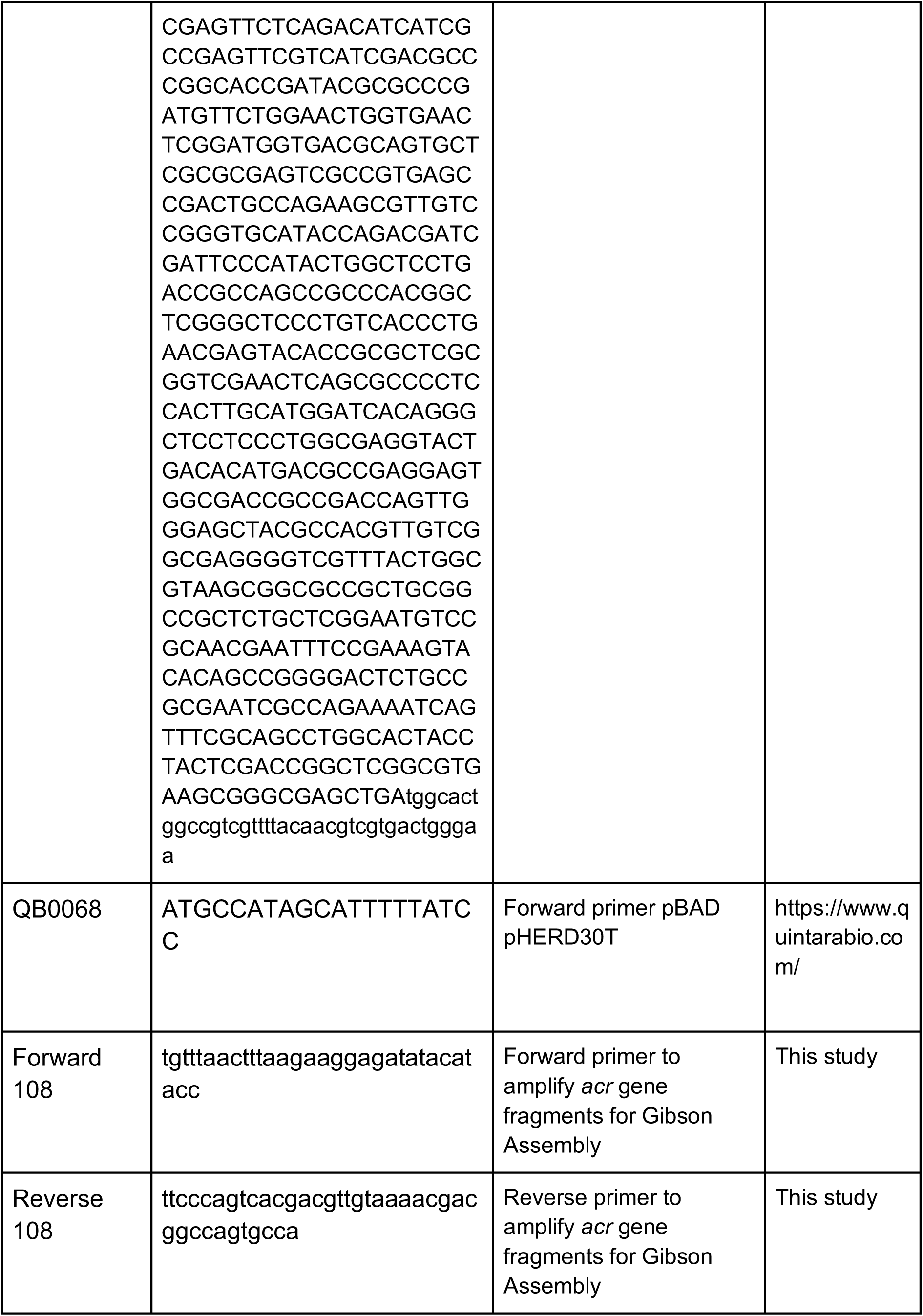
List of oligonucleotides used in this study

**Table S4.**
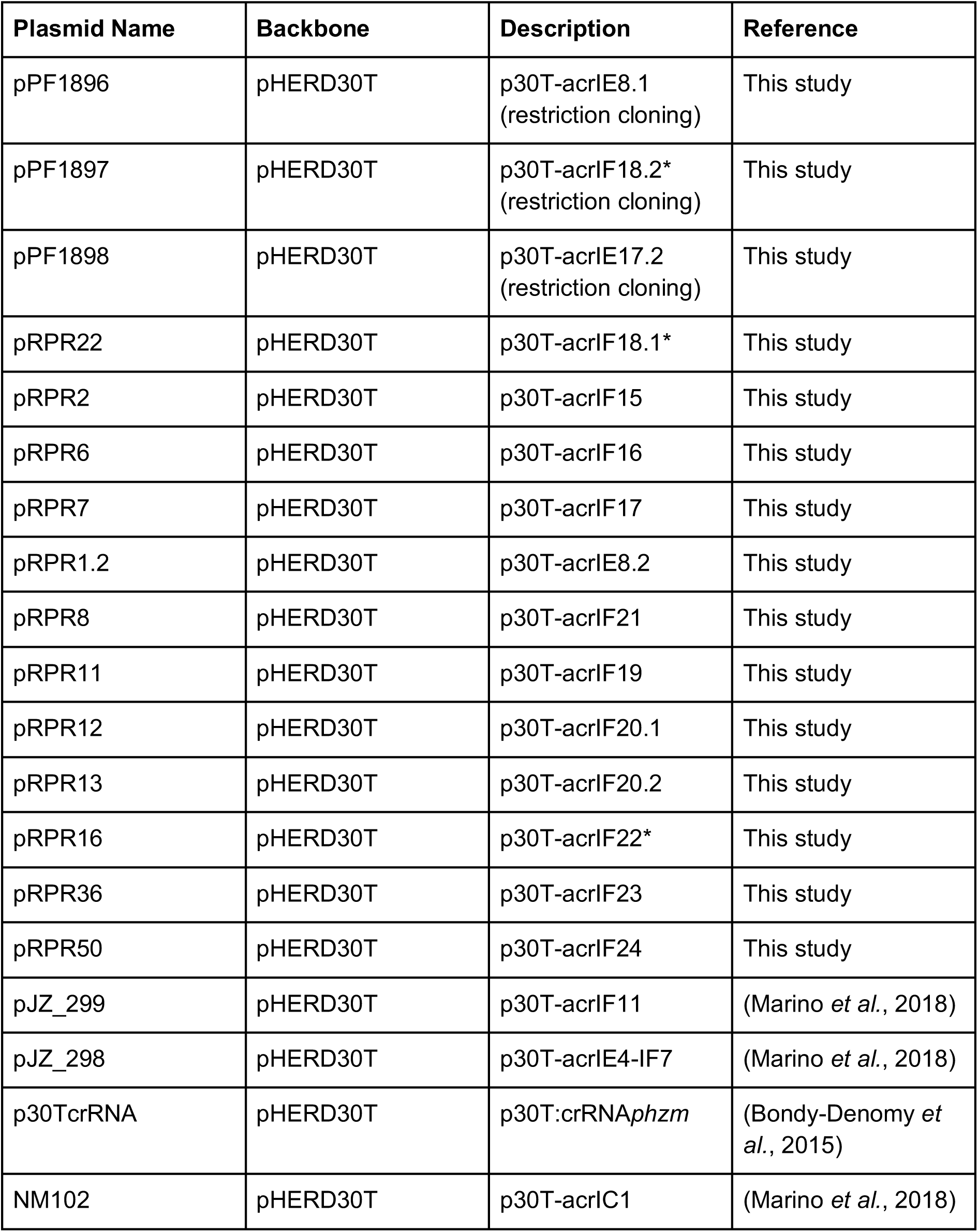
Plasmid constructs used in this study

**Supplementary Table S5.** Software and algorithms

| **Program** | **Source** | **Reference** |
| --- | --- | --- |
| CRISPR/Cas finder | https://crisprcas.i2bc.paris-saclay.fr/ | (Couvin *et al.*, 2018) |
| MegaX | https://www.megasoftware.net/ | (Kumar, Stecher and Tamura, 2016) |
| CRISPRTarget | http://crispr.otago.ac.nz/CRISPRTarget/crispr_analysis.html | (Biswas *et al.*, 2013) |
| Graphpad Prism 6.0 | www.graphpad.com | GraphPad Software, La Jolla California USA. |
| Adobe Illustrator | https://adobe.com/products/illustrator | Adobe Inc., 2019. Adobe Illustrator |
| iTOL | https://itol.embl.de/ | (Letunic and Bork, 2019) |
| HHPred | https://toolkit.tuebingen.mpg.de/tools/hhpred | (Söding, Biegert and Lupas, 2005) |
| Phyre2 | http://www.sbg.bio.ic.ac.uk/~phyre2/html/page.cgi?id=index | (Kelley *et al.*, 2015) |
| HMMER3.0 | https://www.ebi.ac.uk/Tools/hmmer/ | (Potter *et al.*, 2018) |
| Easyfig 2.2.2 | https://mjsull.github.io/Easyfig/ | (Sullivan *et al.*, 2011) |
| IMG/VR | https://img.jgi.doe.gov/vr/ | (Paez-Espino *et al.*, 2017) |
| JPred4 | http://www.compbio.dundee.ac.uk/jpred/ | (Drozdetskiy *et al.*, 2015) |

**Supplementary Figure S1.**
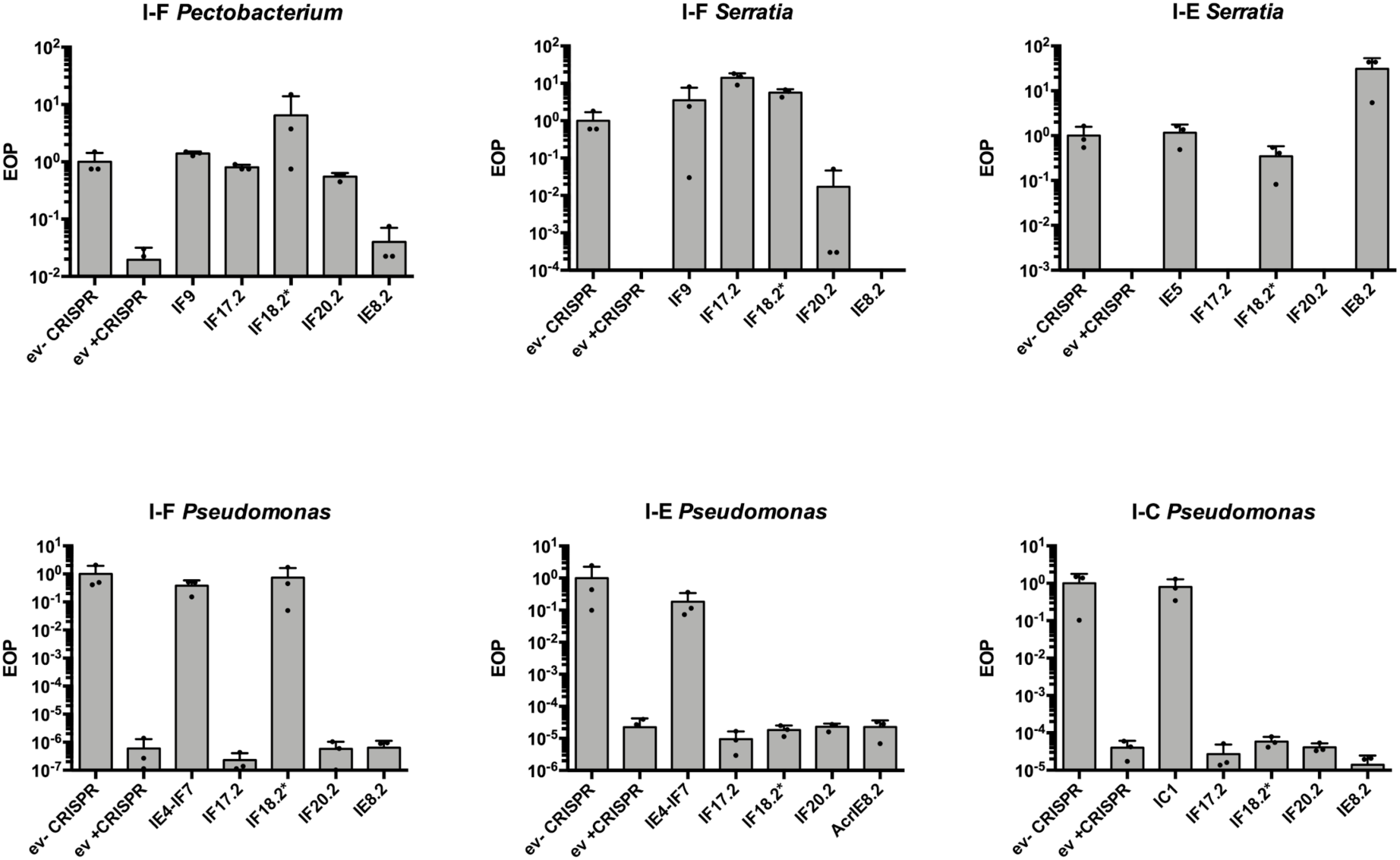
Functional screening of homologs of the acr candidates displayed in figure 2c. Plots show the relative efficiency of plaquing (EOP) of phages in bacterial lawns expressing the different *acr* candidates compared to EOP of the same phage in non-targeting bacterial lawns carrying the empty vector (ev). Asterisk denotes that the Arc in question is a dual I-F and I-E inhibitor.

**Supplementary Figure S2.**
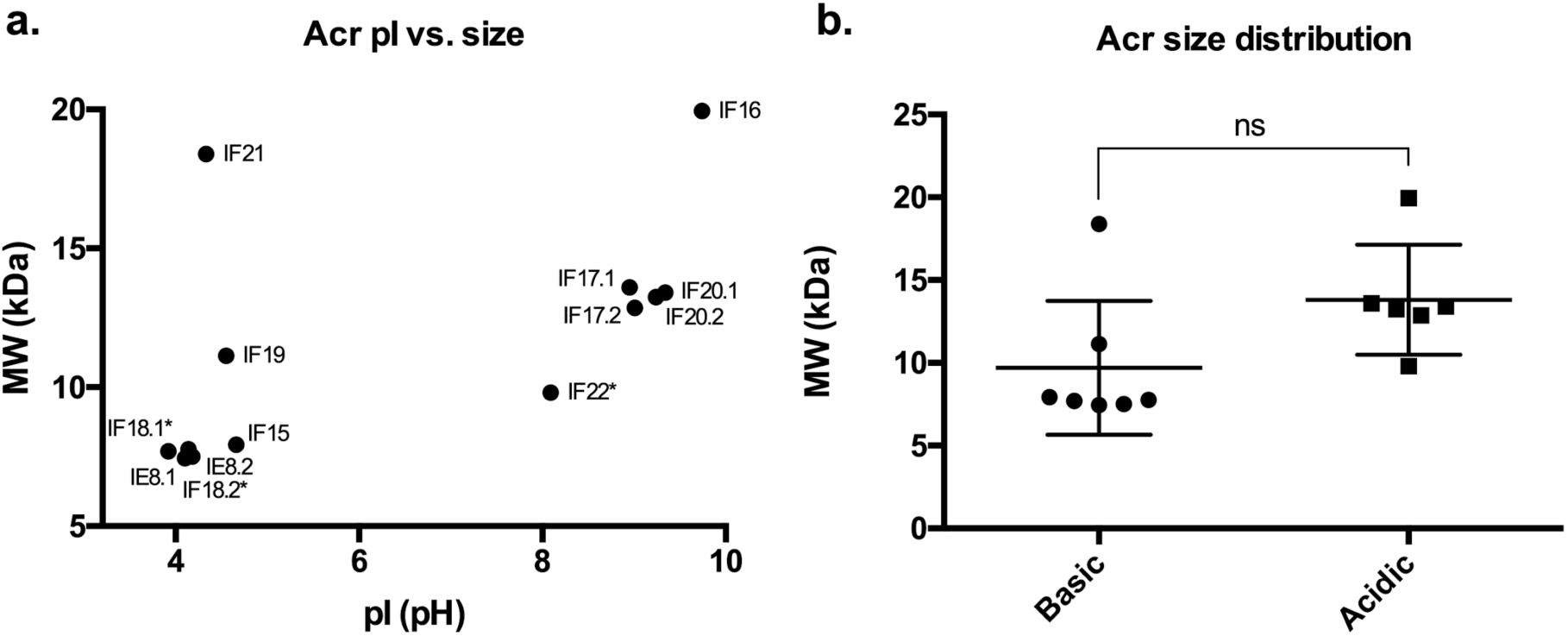
Analysis of the molecular weights (MW) and isoelectric points (pI) of the newly identified I-F and I-E Acrs. **a.** Scatterplot showing the relationship between Acr pI and MW. **b.** Acr distribution for Acrs grouped by acidic (pH < 7) and basic (pH > 7) pIs. No statistical significance (ns) was observed between the basic and acidic Acr groups (unpaired t test; p value = 0.0733). Asterisk denotes that the Arc in question is a dual I-F and I-E inhibitor.

**Supplementary Figure S3.**
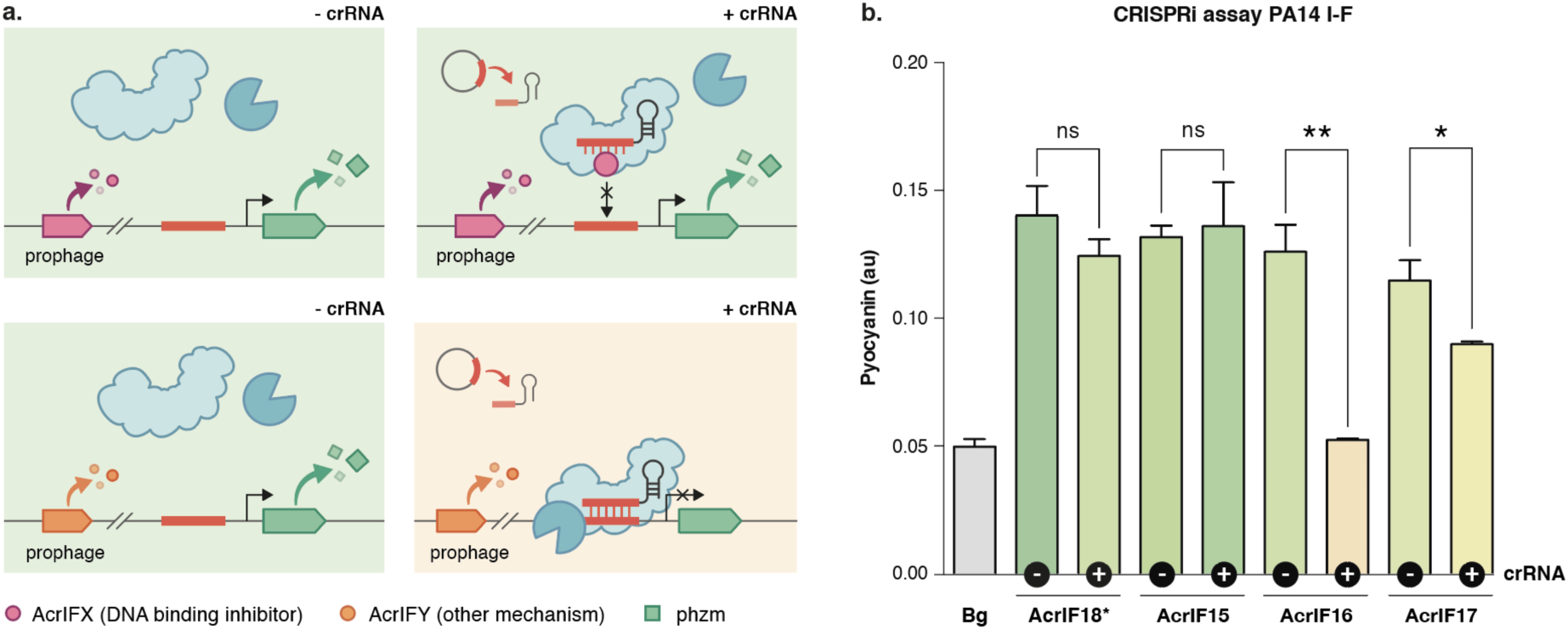
AcrIF18* and AcrIF5 inhibit Cascade target DNA binding. **a.** Schematic of the CRISPRi biochemical assay to test Cascade DNA-binding inhibitory activity. Acrs are expressed in a CRISPRi strain (catalytically dead Cas3 background; dCas3) where the Cascade complex is targeted via a crRNA to the promoter region upstream of *phzM*, a gene involved in the production of the blue-green pigment pyocyanin. Cascade binding effects transcriptional repression of *phzM*, leading to a color change in the culture medium (green to yellow). In the absence of a crRNA, pyocyanin is produced at normal levels (green medium). When a crRNA is provided in the presence of a DNA-binding inhibitor (purple circle), transcription of *phzM* by Cascade is de-repressed (no color change; green). If the Acr does not affect the ability of Cascade to bind it’s DNA target (orange circle), *phzM* is repressed (color change; yellow). **b.** Average pyocyanin production levels for three CRISPRi lysogens expressing a prophage-encoded Acr (AcrIF15, AcrIF16, AcrIF17, and AcrIF18*) in the presence and absence of a crRNA (indicated by the +/− sign in the black circles on the x-axis). “Bg” represents the background pyocyanin detection levels for the assay. Representative colors displayed in the graph have been derived from pictures of the results, see Supplementary Fig S10. Error bars indicate the standard deviation of the mean for three biological replicates.

**Supplementary Figure S4.**
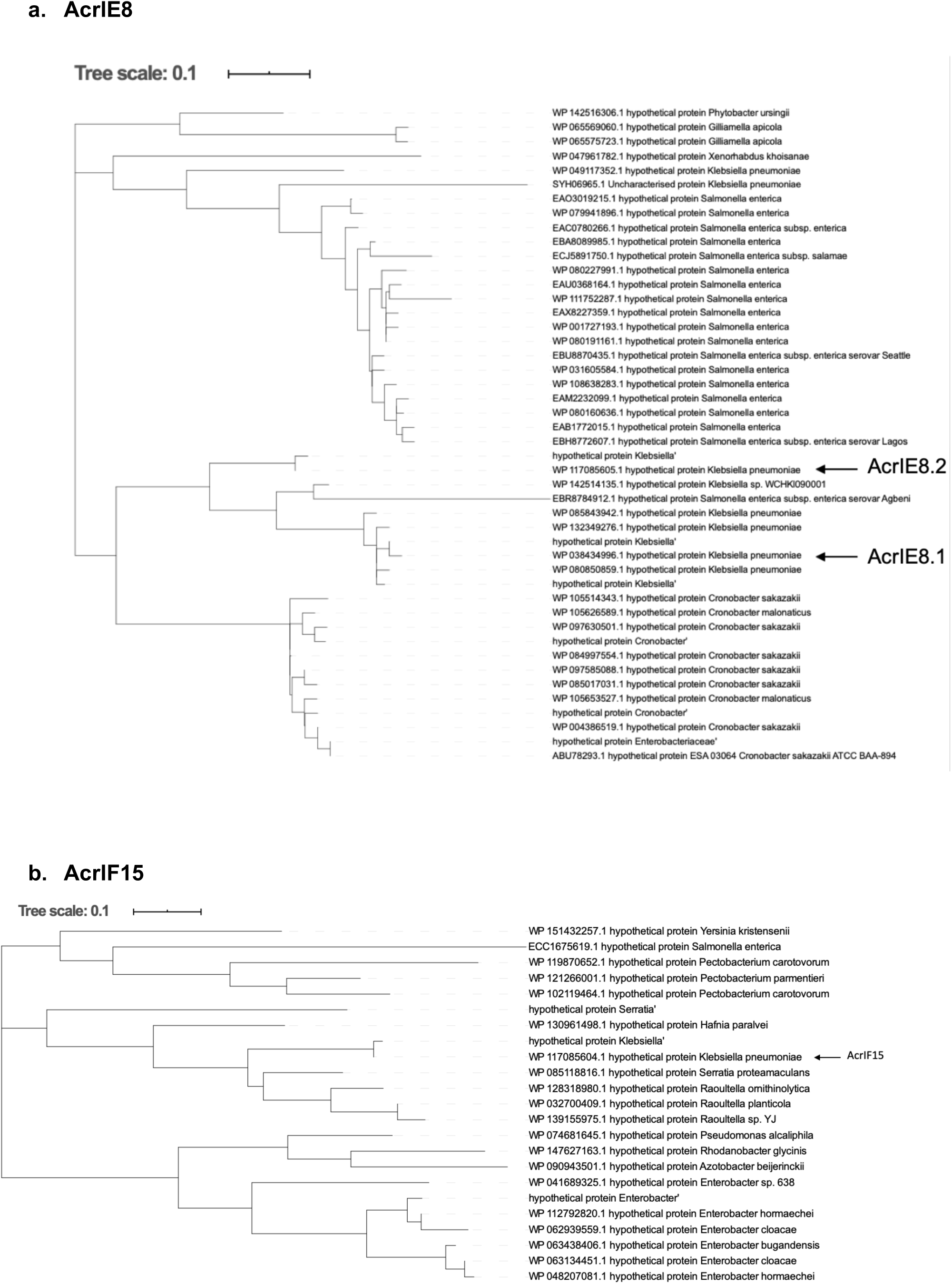
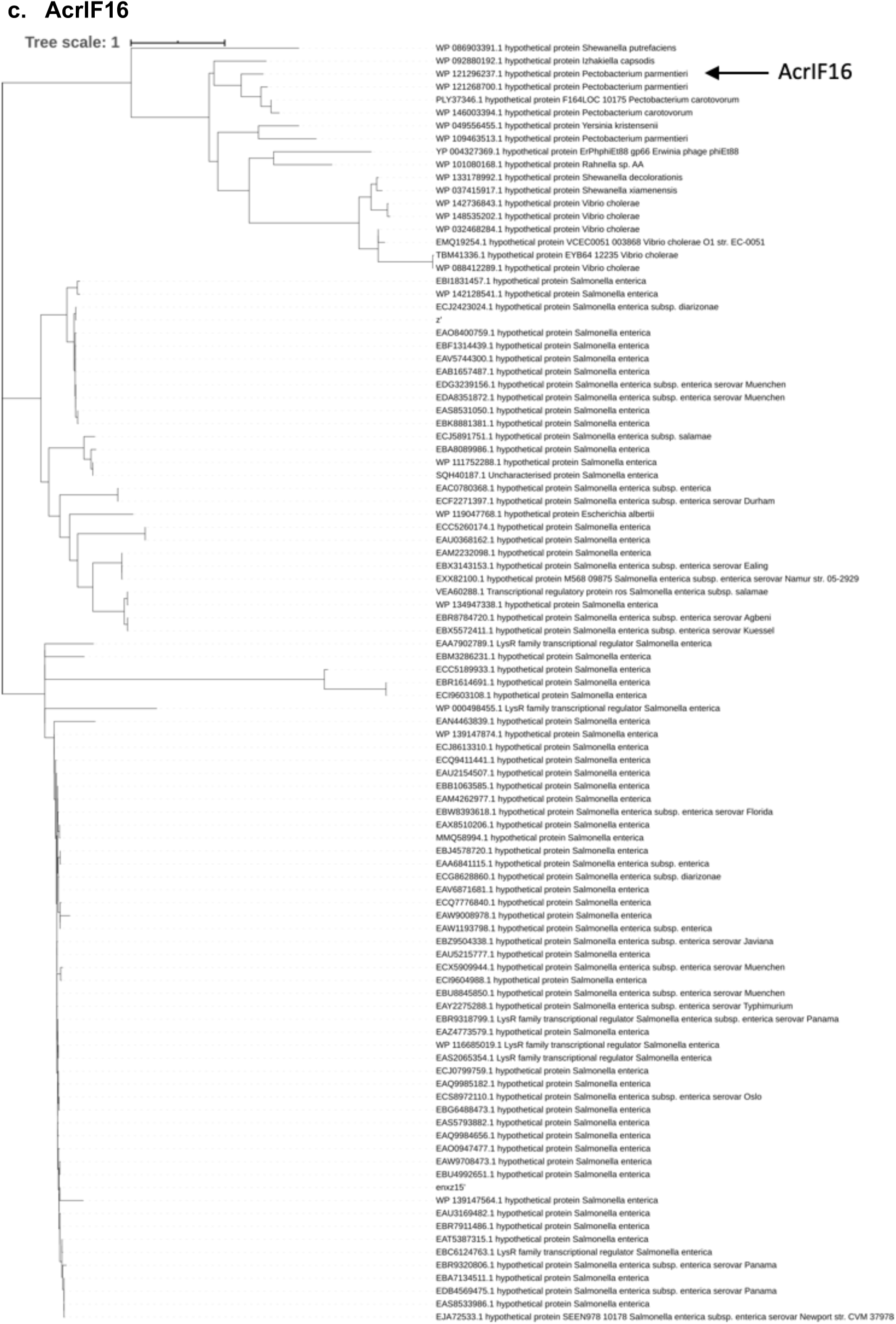
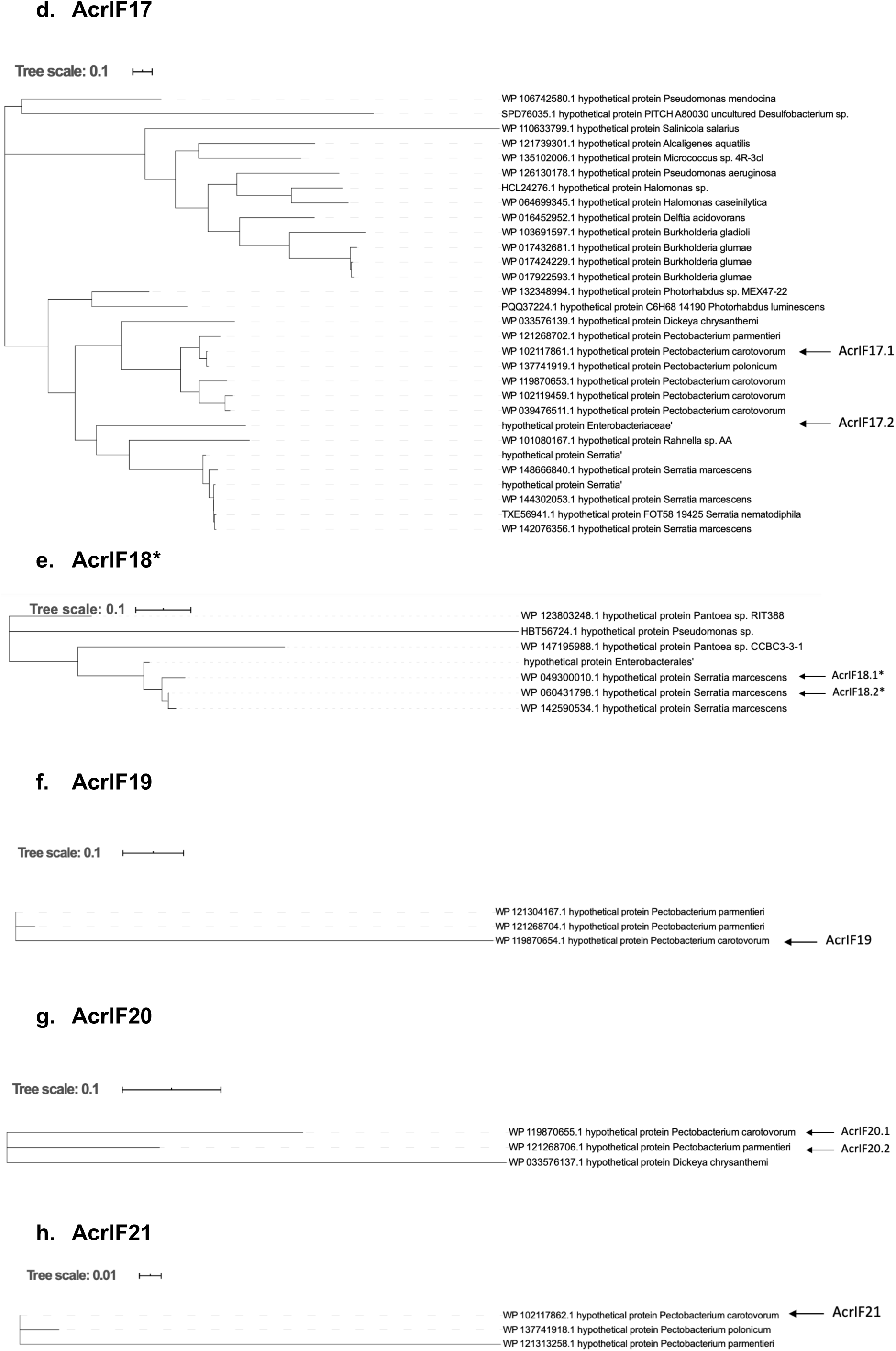
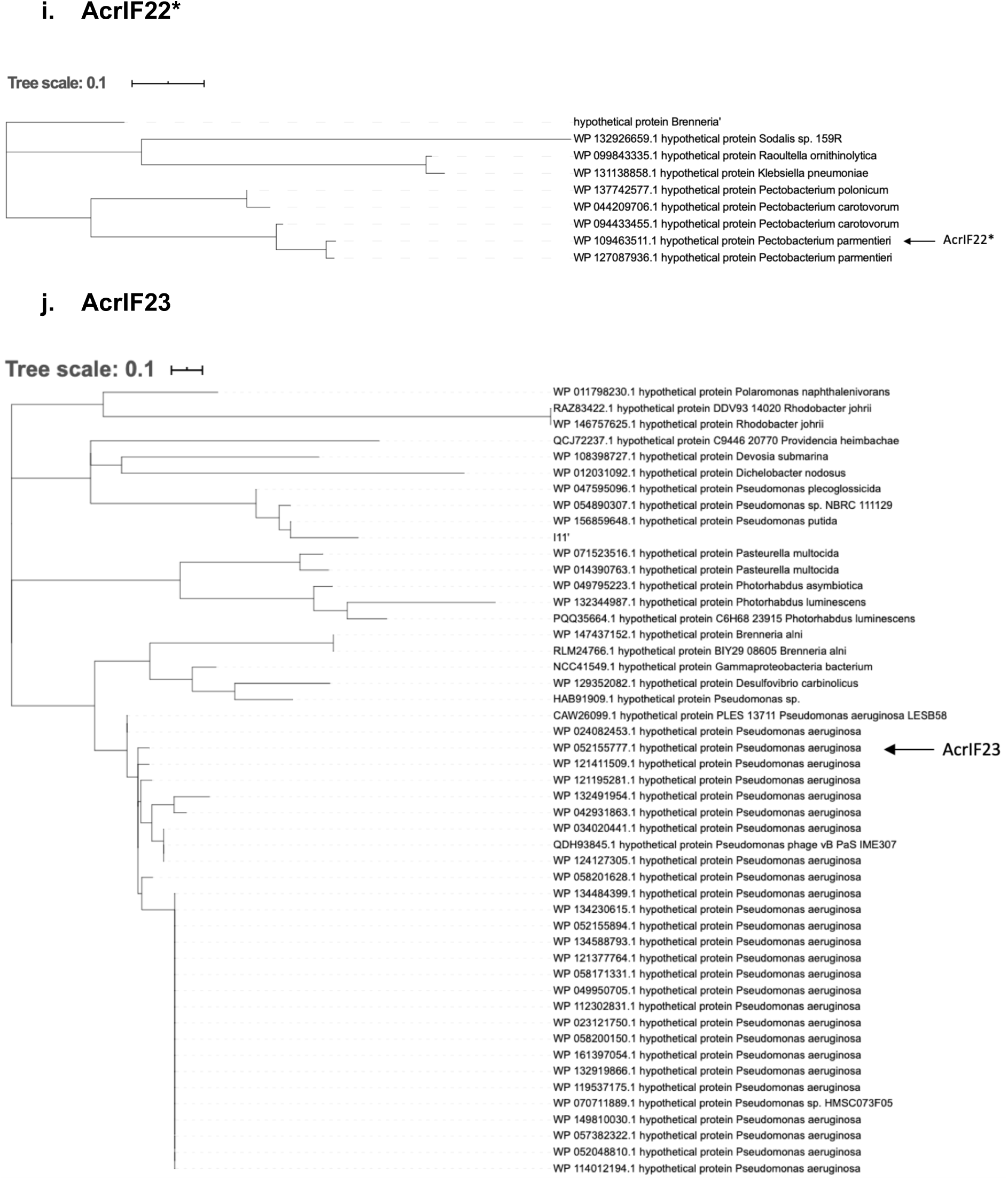
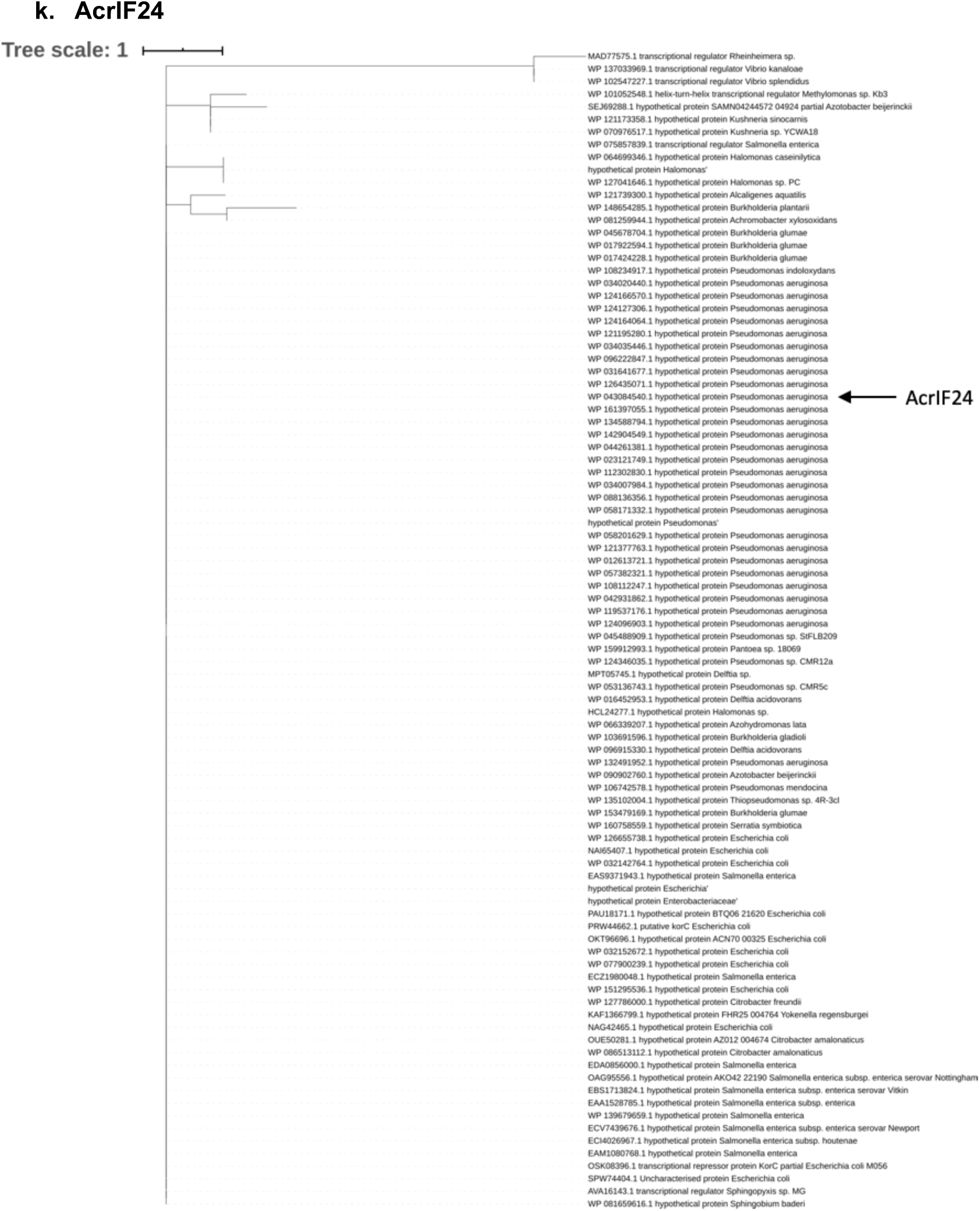
Distribution of the identified Acrs across bacterial taxa. For each Acr family, phylogenetic analyses were performed with the homologs retrieved after PSI-BLAST searches (up to three iterations, only considering hits with e-values <10^−4^ for PSSM generation). Multiple alignments were performed with the MUSCLE software and Maximum Likelihood phylogenetic trees were constructed using MEGA. Branches are labeled with the corresponding protein accession number, followed by the name of the bacterial host species. Arrows indicate the positions in the tree where the Acr homologs tested in this study are found.

**Supplementary Figure S5:**
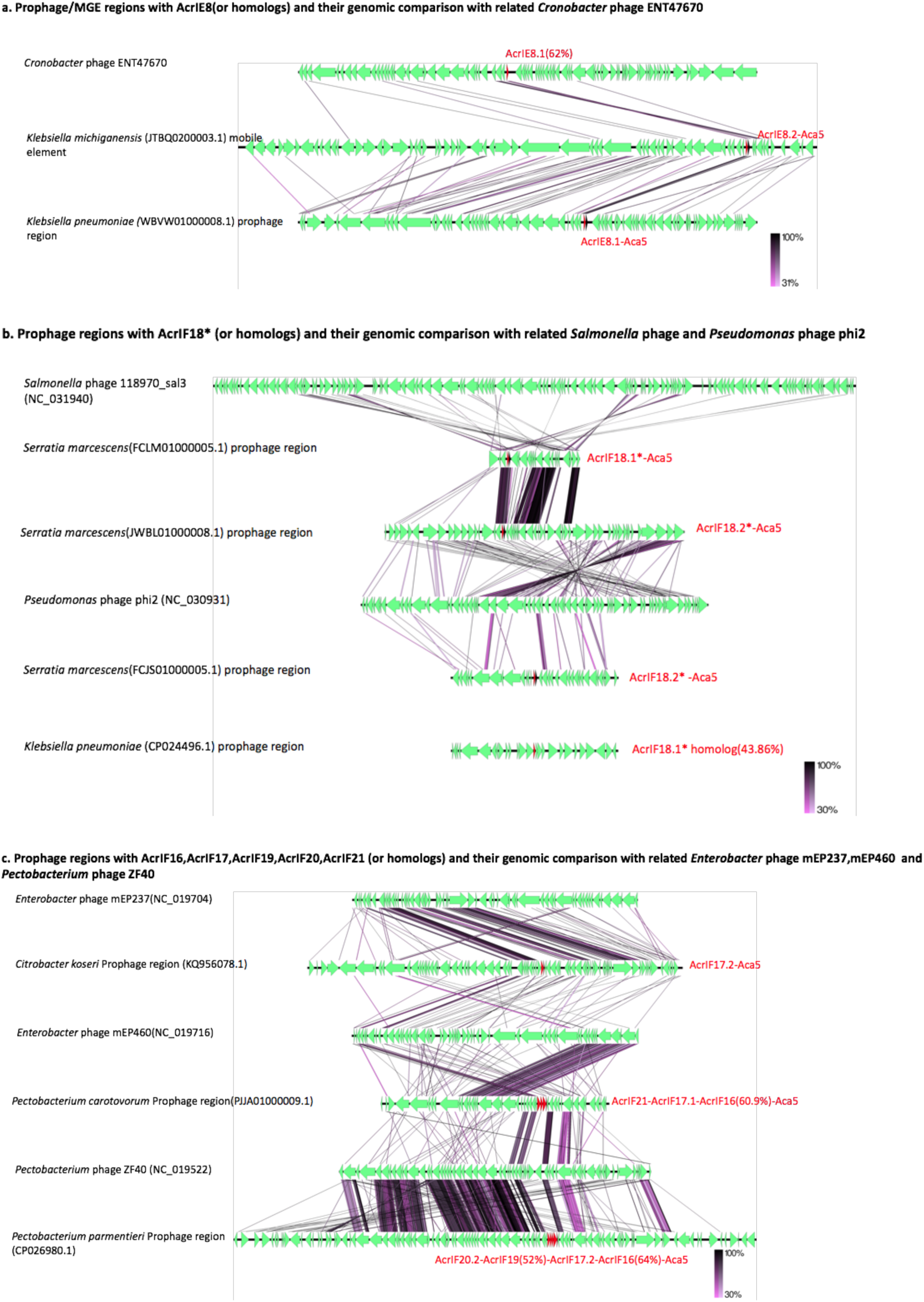
Genome comparison among prophages/MGEs having anti-CRISPRs with related phages: Tblastx has been performed to compare different prophage genomes with their closely related phages (PHASTER best hit). The genomic regions are visualized with EasyFig V2.2.2. All encoded proteins except representative anti-CRISPRs (red) are coloured green. Sequence similarities are shown with gradient (purple to black; between 30% and 100%) straight lines. **(a)** The prophages/MGEs with AcrIE8 share sequence similarity with *Cronobacter* phage ENT47670 genome. All of these genomes have AcrIE8 but the position of *acrs*, neighbouring gene cassettes are variable among different genomes. **(b)** The prophages carrying AcrIF18* share sequence similarity with *Salmonella* phage 118970_sal3 and *Pseudomonas* phage phi2.These phage genomes do not have AcrIF18* and the position of *acrs*, neighbouring gene cassettes are variable among different prophage genomes. **(c)** The anti-CRISPRs AcrIF16, AcrIF17, AcrIF19, AcrIF20, AcrIF21 and the homologs of these Acrs are mostly found to be clustered in the same prophage regions. These regions have sequence similarity with *Enterobacter* phage mEP460, mEP237, and *Pectobacterium* phage ZF40. The phage genomes do not have these anti-CRISPRs, and the position of *acrs*, neighbouring gene cassettes are variable among different prophage genomes. A detailed analysis with *Pectobacterium* phage ZF40 is described in the text and Figure 3.

**Supplementary Figure S6.**
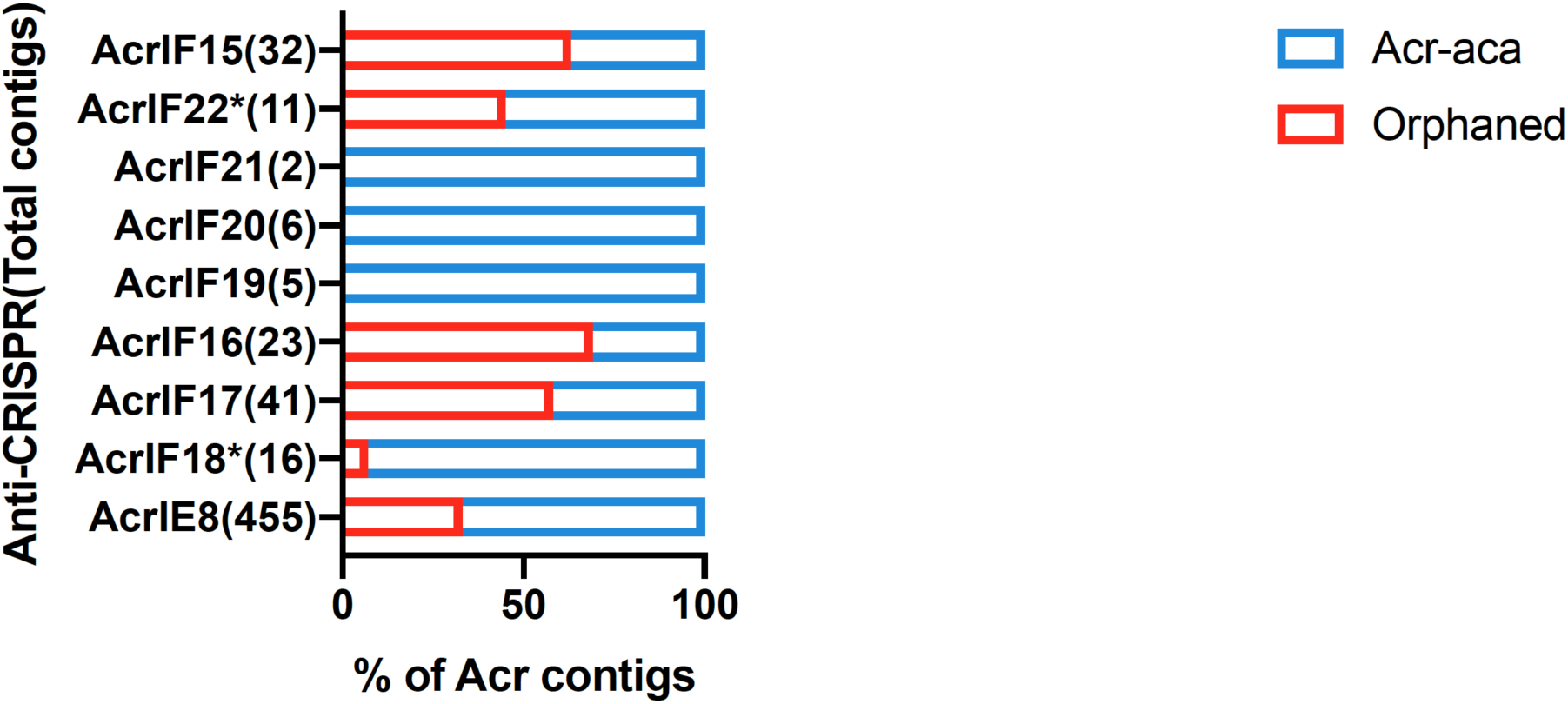
Proportion of newly identified acrs associated with a known *aca* gene. Homology search of the different acrs presented in this work against all Refseq bacterial contigs (total n = 18654021) revealed a significant number of orphaned Acrs in the bacterial genomes.

**Supplementary Figure S7.**
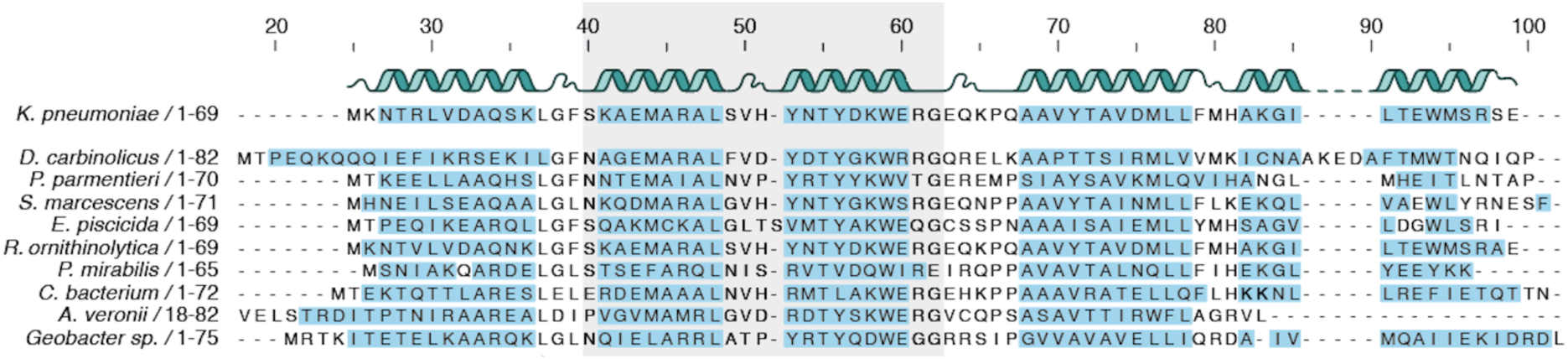
Secondary structure prediction for the diverse Aca9 homologs displayed in the alignment form Figure 4b. Predicted secondary structure alpha-helix regions are illustrated as ribbons for the *K. pneumoniae* homolog (top) and blue shadings indicate the corresponding predictions across the different orthologs. The predicted HTH DNA binding domain is highlighted with a gray shaded area.

**Supplementary Figure S8.**
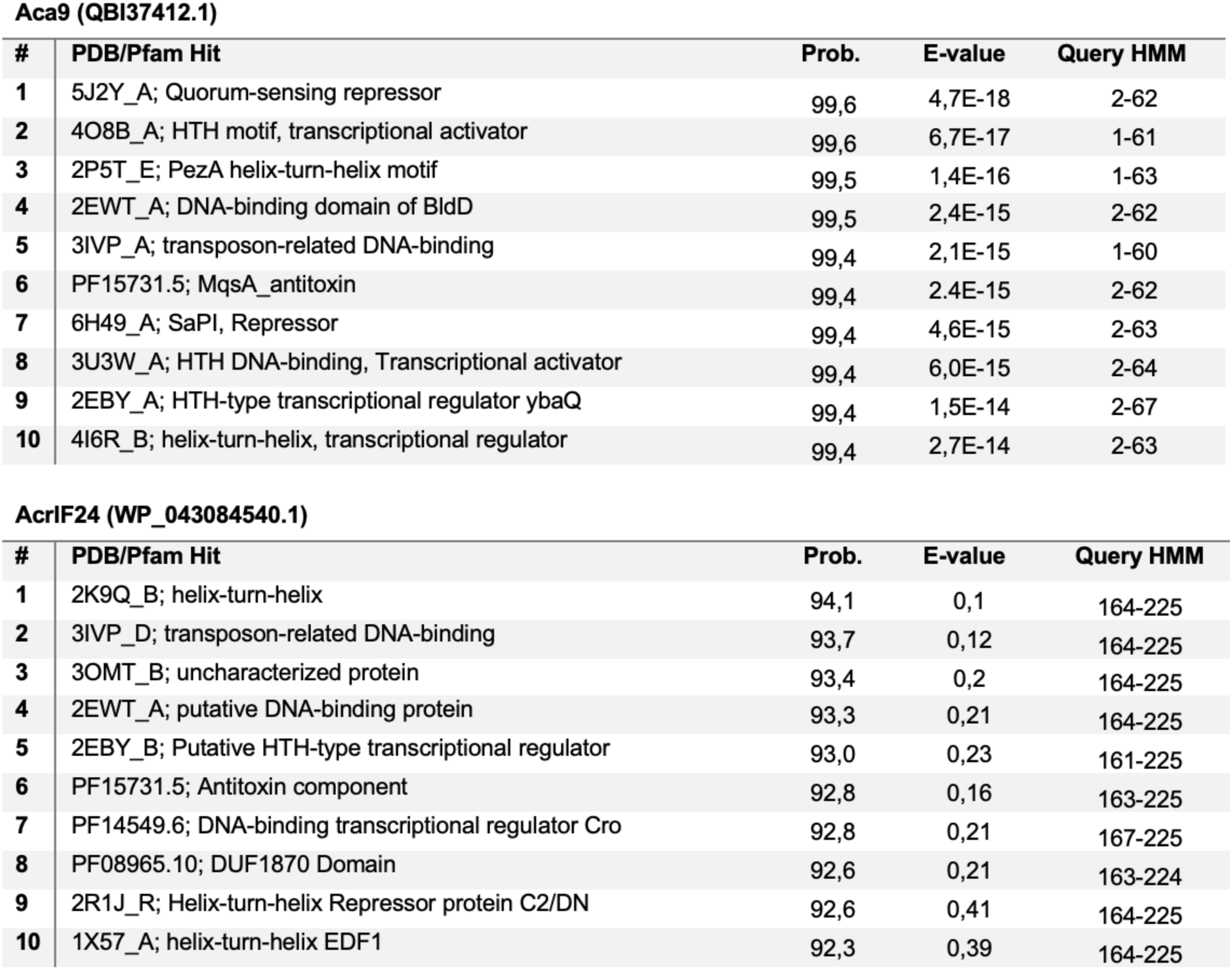
Summary of the HHPred analyses performed for Aca9 and AcrIF24. The top ten PDB/Pfam protein hits and associated domains ranked according to the HHPred probability score are listed, together with the corresponding e-values and matching amino acid positions in the protein query (query HMM).

**Supplementary Figure S9.**
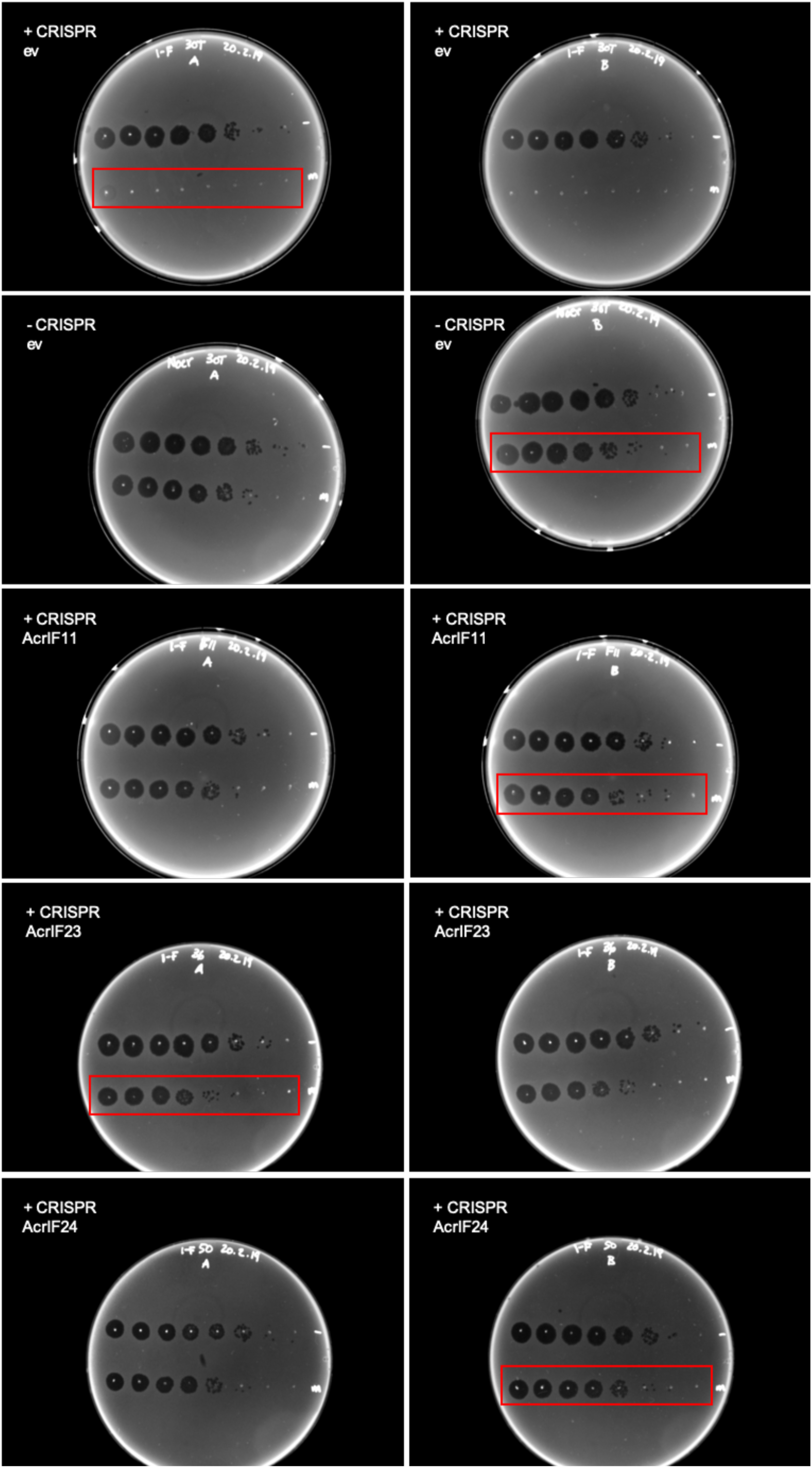
Image manipulation was performed to enhance the visibility of the phage titering results shown in “Figure 4d”. The raw, un-cropped images are shown here, highlighting (red rectangles) the portion of the original photographs displayed in Figure 4d. The plate images show the results of the functional validation experiments for AcrIF23 and AcrIF24. Briefly, PA14 lawns (+CRISPR) were spotted with 10-fold dilutions of CRISPR insensitive DMS3 phage (top dilution row) and CRISPR sensitive DMS3m phage (bottom dilution row), in the presence of the empty vector pHERD30T (ev), AcrIF11 (previously validated Acr; positive control for inhibition), AcrIF23, and AcrIF24. Lawns of a non-targeting PA14 strain (-CRISPR) were transformed with the empty vector (ev) as a negative control for DMS3m targeting.

**Supplementary Figure S10.**
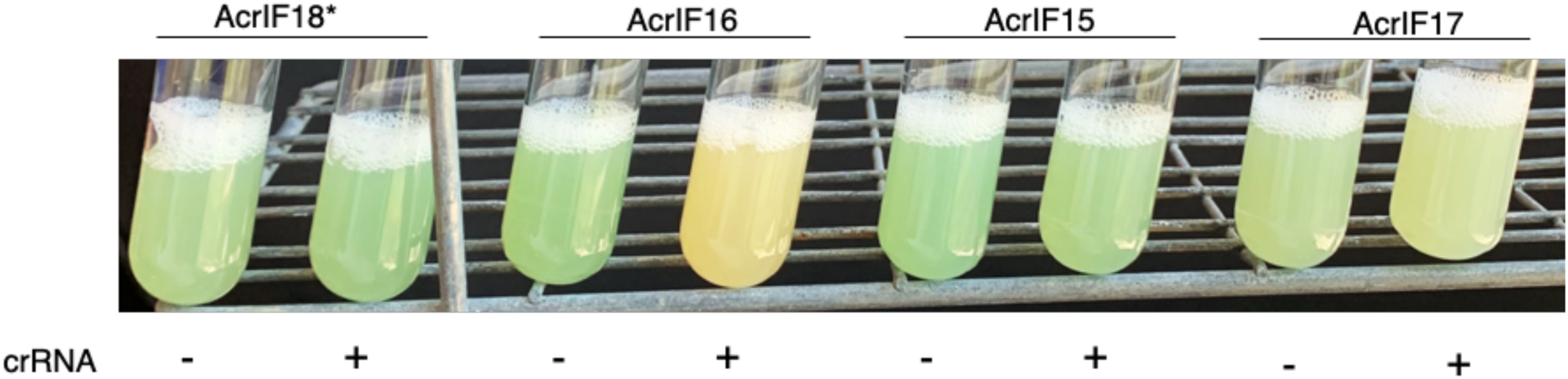
CRISPRi-based repression of pyocyanin production. Assay results showing a representative picture of the observed change in color observed for the expression of AcrIF18*, AcrIF16, AcrIF15 and AcrIF17, in the presence (+) or absence (-) of a crRNA targeting the *phzM* gene, in a delta Cas3 PA14 genomic background.

## Supplementary Datasheets

**Supplementary Datasheet S1. Information summary for the identified Acrs.** Information includes: the accession numbers for the different Acrs (and homologs) tested in this work, predicted isoelectric points (pI) for the proteins, amino acid (aa) sequence lengths, molecular weights (MW), positions in the genomes of the bacteria where *acrs* are found, positions and (sub)types of detected CRISPR arrays and Cas operons in the bacterial host genomes, presence/absence of self-targeting within the host genomes, positions of the identified prophage regions found within the genomes of bacteria encoding a validated Acrs.

**Supplementary Datasheet S2. List and analysis of self-targeting spacers**. The following information is provided: the spacer number that self-targeting spacers occupy in the array of origin and corresponding CRISPR-Cas subtype, the genomic position (bp) where self targeting occurs, the sequence of the self-targeting spacer, number of mismatches, predicted PAMs, whether a phage is predicted within the predicted self-targeted region, the ORF and predicted protein-coding genes targeted (when possible).

**Supplementary Datasheet S3. CRISPR-Cas sequence identity comparison calculations used to make Figure 2b.** The results from BLATSp comparisons between the Cas orthologs of the different model CRISPR-Cas systems tested and the selected endogenous CRISPR-Cas systems (hosts from which the selected *acrs* originate) is shown. An average score of the percentage sequence identities was calculated for all one-to-one comparisons.

**Supplementary Datasheet S4. Analysis of the genomic contexts and MGE origins of the homologs of the validated Acrs presented in Figure 2.** The datasheet includes a list of accession numbers and protein sequences for all identified homologs of AcrIE8, AcrIF15, AcrIF16, AcrIF17, AcrIF18*, AcrIF19, AcrIF16, AcrIF20, AcrIF21, AcrIF22* that were used to build the phylogenetic trees of the Acr families. MGE-related genes used to determine the genetic context of each Acr homolog are listed. A summary count of the homologs found in phage, plasmid, and unknown genomic contexts is provided for each Acr.

**Supplementary Datasheet S5.** List of Aca9 homologs used to build the phylogenetic tree in Figure 4. MGE-related genes used to determine the genetic context of each homolog are listed.

**Supplementary Datasheet S6.** List of putative antibiotic resistance genes identified on the *Klebsiella pneumoniae* plasmid shown on Figure 5a. BLAST searches were performed against the Comprehensive Antibiotic Resistance Database (CARD). Positions for the location of genes in the plasmid genome, the e-value for the matches, and a short description of the gene function are indicated.

**Supplementary Datasheet S7.** Lists of accession numbers and protein sequences for the homologs used to build the Aca5 phylogenetic tree in Figure 1a.

**Supplementary Datasheet S8.** Lists of accession numbers and protein sequences for the homologs used to build the AcrIF23 and AcrIF24 phylogenetic trees in Supplementary Figure S3.

**Supplementary Datasheet S9. Raw data for the acr candidate validation experiments.** Plaque forming unit count data and efficiency of plaquing calculations presented in Figure 2c.

**Supplementary Datasheet S10. HMM search results for the Acrs presented in this study.** Top hits obtained from searches against pfam are shown. The probability, e-value and matching amino acid positions in the protein query (query HMM) are indicated.

**Supplementary Datasheet S11. Pyocyanin repression assay: raw data.** Pyocyanin was extracted from the overnight cultures and quantified by measuring absorbance at 520 nm.

## Acknowledgements

We thank members of our laboratories for helpful discussions. We are thankful to Marina Pinilla for her assistance in the creative design of figues. Thanks to Lucia Malone for providing JS26-targeting *Serratia* strains, Nils Birkholz for demonstrating plaque assays to S.S and Te-yuan Chyou for providing modified HMMER script. Financial support for R.P-R. was provided by the Innovation Fund Denmark, Trojan Horse Project [#5157-00005B], the Independent Research Fund Denmark, InTrans Project [#8022-00322B], and the JPI-AMR DARWIN Project [#7044-00004B]. P.C.F and R.D.F were supported in part by the Marsden Fund from the Royal Society of New Zealand and P.C.F. by the Bio-protection Research Centre (Tertiary Education Commission, NZ). S.S was supported by a University of Otago Doctoral Scholarship. N.D.M. was supported by NIH F32GM133127. J.B.-D. And Bondy-Denomy lab was supported by the UCSF Program for Breakthrough Biomedical Research funded in part by the Sandler Foundation, the Searle Fellowship, the Vallee Foundation, the Innovative Genomics Institute, an NIH Director’s Early Independence Award DP5-OD021344, R01GM127489, and by DARPA HR0011-17-2-0043.

## References

Arndt, D. et al. (2016) ‘PHASTER: a better, faster version of the PHAST phage search tool’, Nucleic acids research, 44(W1), pp. W16–21.

Athukoralage, J. S. et al. (2020) ‘An anti-CRISPR viral ring nuclease subverts type III CRISPR immunity’, Nature, 577(7791), pp. 572–575.

Bagdasarian, M. et al. (1986) ‘An inhibitor of SOS induction, specified by a plasmid locus in Escherichia coli’, Proceedings of the National Academy of Sciences of the United States of America, 83(15), pp. 5723–5726.

Bell, K. S. et al. (2004) ‘Genome sequence of the enterobacterial phytopathogen Erwinia carotovora subsp. atroseptica and characterization of virulence factors’, Proceedings of the National Academy of Sciences of the United States of America, 101(30), pp. 11105–11110.

Bernheim, A. and Sorek, R. (2019) ‘The pan-immune system of bacteria: antiviral defence as a community resource’, Nature reviews. Microbiology. doi: 10.1038/s41579-019-0278-2.

Bhoobalan-Chitty, Y. et al. (2019) ‘Inhibition of Type III CRISPR-Cas Immunity by an Archaeal Virus-Encoded Anti-CRISPR Protein’, Cell, pp. 448–458.e11. doi: 10.1016/j.cell.2019.09.003.

Birkholz, N. et al. (2019) ‘The autoregulator Aca2 mediates anti-CRISPR repression’, Nucleic acids research, 47(18), pp. 9658–9665.

Biswas, A. et al. (2013) ‘CRISPRTarget: bioinformatic prediction and analysis of crRNA targets’, RNA biology, 10(5), pp. 817–827.

Blower, T. R. et al. (2012) ‘Viral Evasion of a Bacterial Suicide System by RNA–Based Molecular Mimicry Enables Infectious Altruism’, PLoS Genetics, p. e1003023. doi: 10.1371/journal.pgen.1003023.

Bondy-Denomy et al. (2013) ‘Bacteriophage genes that inactivate the CRISPR/Cas bacterial immune system’, Nature, 493(7432), pp. 429–432.

Bondy-Denomy, J. et al. (2015) ‘Multiple mechanisms for CRISPR-Cas inhibition by anti-CRISPR proteins’, Nature, 526(7571), pp. 136–139.

Borges, A. L. et al. (2018) ‘Bacteriophage Cooperation Suppresses CRISPR-Cas3 and Cas9 Immunity’, Cell, pp. 917–925.e10. doi: 10.1016/j.cell.2018.06.013.

Borges, A. L., Davidson, A. R. and Bondy-Denomy, J. (2018) ‘The Discovery, Mechanisms, and Evolutionary Impact of Anti-CRISPRs’, Annual review of virology, 4(1), pp. 37–59.

Blower, T. R., Evans, T. J., Przybilski, R., Fineran, P. C., & Salmond, G. P. (2012). Viral evasion of a bacterial suicide system by RNA–based molecular mimicry enables infectious altruism. PLoS genetics, 8(10).

Cady, K. C. et al. (2012) ‘The CRISPR/Cas adaptive immune system of Pseudomonas aeruginosa mediates resistance to naturally occurring and engineered phages’, Journal of bacteriology, 194(21), pp. 5728–5738.

Chowdhury, S. et al. (2017) ‘Structure Reveals Mechanisms of Viral Suppressors that Intercept a CRISPR RNA-Guided Surveillance Complex’, Cell, 169(1), pp. 47–57.e11.

Cohen, D. et al. (2019) ‘Cyclic GMP–AMP signalling protects bacteria against viral infection’, Nature, pp. 691–695. doi: 10.1038/s41586-019-1605-5.

Couvin, D. et al. (2018) ‘CRISPRCasFinder, an update of CRISRFinder, includes a portable version, enhanced performance and integrates search for Cas proteins’, Nucleic acids research, 46(W1), pp. W246–W251.

Davidson, A. R. et al. (2020) ‘Anti-CRISPRs: Protein Inhibitors of CRISPR-Cas Systems’, Annual review of biochemistry. doi: 10.1146/annurev-biochem-011420-111224.

Dong, D. et al. (2017) ‘Structural basis of CRISPR–SpyCas9 inhibition by an anti-CRISPR protein’, Nature, pp. 436–439. doi: 10.1038/nature22377.

Dorman, C. J. (2014) ‘H-NS-like nucleoid-associated proteins, mobile genetic elements and horizontal gene transfer in bacteria’, Plasmid, 75, pp. 1–11.

Dorman, C. J. and Ní Bhriain, N. (2020) ‘CRISPR-Cas, DNA Supercoiling, and Nucleoid-Associated Proteins’, Trends in microbiology, 28(1), pp. 19–27.

Doron, S. et al. (2018) ‘Systematic discovery of antiphage defense systems in the microbial pangenome’, Science, 359(6379). doi: 10.1126/science.aar4120.

Drozdetskiy, A. et al. (2015) ‘JPred4: a protein secondary structure prediction server’, Nucleic Acids Research, pp. W389–W394. doi: 10.1093/nar/gkv332.

Edgar, R. C. (2004) ‘MUSCLE: multiple sequence alignment with high accuracy and high throughput’, Nucleic acids research, 32(5), pp. 1792–1797.

Fineran, P. C. et al. (2014) ‘Degenerate target sites mediate rapid primed CRISPR adaptation’, Proceedings of the National Academy of Sciences of the United States of America, 111(16), pp. E1629–38.

Finn, R.D., Clements, J. and Eddy, S.R., 2011. “HMMER web server: interactive sequence similarity searching.” Nucleic acids research, 39: 29–37.

Forsberg, K. J. et al. (2019) ‘Functional metagenomics-guided discovery of potent Cas9 inhibitors in the human microbiome’, eLife, 8. doi: 10.7554/eLife.46540.

Gleditzsch, D. et al. (2019) ‘PAM identification by CRISPR-Cas effector complexes: diversified mechanisms and structures’, RNA biology, 16(4), pp. 504–517.

Guglielmini, J. et al. (2011) ‘The Repertoire of ICE in Prokaryotes Underscores the Unity, Diversity, and Ubiquity of Conjugation’, PLoS Genetics, p. e1002222. doi: 10.1371/journal.pgen.1002222.

Günthert, U. and Reiners, L. (1987) ‘Bacillus subtilis phage SPR codes for a DNA methyltransferase with triple sequence specificity’, Nucleic acids research, 15(9), pp. 3689–3702.

Guo, T. W. et al. (2017) ‘Cryo-EM Structures Reveal Mechanism and Inhibition of DNA Targeting by a CRISPR-Cas Surveillance Complex’, Cell, 171(2), pp. 414–426.e12.

Hampton, H. G., Watson, B. N. J. and Fineran, P. C. (2020) ‘The arms race between bacteria and their phage foes’, Nature, 577(7790), pp. 327–336.

Hille, F. et al. (2018) ‘The Biology of CRISPR-Cas: Backward and Forward’, Cell, 172(6), pp. 1239–1259.

Hynes, A. P. et al. (2018) ‘Widespread anti-CRISPR proteins in virulent bacteriophages inhibit a range of Cas9 proteins’, Nature communications. doi: 10.1038/s41467-018-05092-w.

Isaev, A. et al. (2020) ‘Phage T7 DNA mimic protein Ocr is a potent inhibitor of BREX defence’, Nucleic acids research. doi: 10.1093/nar/gkaa290.

Jackson, S. A. et al. (2019) ‘Imprecise Spacer Acquisition Generates CRISPR-Cas Immune Diversity through Primed Adaptation’, Cell host & microbe, 25(2), pp. 250–260.e4.

Jackson, S.A., Birkholz, N., Malone, L.M. and Fineran, P.C., 2019. Imprecise spacer acquisition generates CRISPR-Cas immune diversity through primed adaptation. Cell host & microbe, 25(2), pp.250–260.

Jiang, F. et al. (2019) ‘Temperature-Responsive Competitive Inhibition of CRISPR-Cas9’, Molecular cell, 73(3), pp. 601–610.e5.

Kelley, L. A. et al. (2015) ‘The Phyre2 web portal for protein modeling, prediction and analysis’, Nature Protocols, pp. 845–858. doi: 10.1038/nprot.2015.053.

Kumar, S., Stecher, G. and Tamura, K. (2016) ‘MEGA7: Molecular Evolutionary Genetics Analysis Version 7.0 for Bigger Datasets’, Molecular Biology and Evolution, pp. 1870–1874. doi: 10.1093/molbev/msw054.

Lee, J. et al. (2018) ‘Potent Cas9 Inhibition in Bacterial and Human Cells by AcrIIC4 and AcrIIC5 Anti-CRISPR Proteins’, mBio, 9(6). doi: 10.1128/mBio.02321-18.

Letunic, I. and Bork, P. (2019) ‘Interactive Tree Of Life (iTOL) v4: recent updates and new developments’, Nucleic acids research, 47(W1), pp. W256–W259.

Liu, L. et al. (2019) ‘Phage AcrIIA2 DNA Mimicry: Structural Basis of the CRISPR and Anti-CRISPR Arms Race’, Molecular cell, 73(3), pp. 611–620.e3.

Mahendra, C. et al. (2020) ‘Broad-spectrum anti-CRISPR proteins facilitate horizontal gene transfer’, Nature microbiology, 5(4), pp. 620–629.

Makarova, K. S. et al. (2011) ‘Defense islands in bacterial and archaeal genomes and prediction of novel defense systems’, Journal of bacteriology, 193(21), pp. 6039–6056.

Makarova, K. S. et al. (2015) ‘An updated evolutionary classification of CRISPR–Cas systems’, Nature reviews. Microbiology. Nature Publishing Group, a division of Macmillan Publishers Limited. All Rights Reserved., 13, p. 722.

Makarova, K. S. et al. (2020) ‘Evolutionary classification of CRISPR–Cas systems: a burst of class 2 and derived variants’, Nature Reviews Microbiology, 18, pp. 67–83.

Marino, N. D. et al. (2018) ‘Discovery of widespread type I and type V CRISPR-Cas inhibitors’, Science, 362(6411), pp. 240–242.

Marino, N. D. et al. (2020) ‘Anti-CRISPR protein applications: natural brakes for CRISPR-Cas technologies’, Nature Methods. doi: 10.1038/s41592-020-0771-6.

Marshall, R. et al. (2018) ‘Rapid and Scalable Characterization of CRISPR Technologies Using an E. coli Cell-Free Transcription-Translation System’, Molecular Cell, pp. 146–157.e3. doi: 10.1016/j.molcel.2017.12.007.

McArthur, A. G. et al. (2013) ‘The comprehensive antibiotic resistance database’, Antimicrobial agents and chemotherapy, 57(7), pp. 3348–3357.

Murphy, J. et al. (2013) ‘Bacteriophage orphan DNA methyltransferases: insights from their bacterial origin, function, and occurrence’, Applied and environmental microbiology, 79(24), pp. 7547–7555.

Norman, A., Hansen, L. H. and Sørensen, S. J. (2009) ‘Conjugative plasmids: vessels of the communal gene pool’, Philosophical transactions of the Royal Society of London. Series B, Biological sciences, 364(1527), pp. 2275–2289.

Osuna, B. A., Karambelkar, S., Mahendra, C., Sarbach, A., et al. (2020) ‘Critical Anti-CRISPR Locus Repression by a Bi-functional Cas9 Inhibitor’, Cell host & microbe. doi: 10.1016/j.chom.2020.04.002.

Osuna, B. A., Karambelkar, S., Mahendra, C., Christie, K. A., et al. (2020) ‘Listeria phages induce Cas9 degradation to protect lysogenic genomes’, Cell Host and Microbe. doi: 10.1101/787200.

Paez-Espino, D. et al. (2017) ‘IMG/VR: a database of cultured and uncultured DNA Viruses and retroviruses’, Nucleic acids research, 45(D1), pp. D457–D465.

Pawluk, A. et al. (2014) ‘A New Group of Phage Anti-CRISPR Genes Inhibits the Type I-E CRISPR-Cas System of Pseudomonas aeruginosa’, mBio. doi: 10.1128/mbio.00896-14.

Pawluk, A., Staals, R. H. J., et al. (2016) ‘Inactivation of CRISPR-Cas systems by anti-CRISPR proteins in diverse bacterial species’, Nature microbiology, 1(8), p. 16085.

Pawluk, A., Amrani, N., et al. (2016) ‘Naturally Occurring Off-Switches for CRISPR-Cas9’, Cell, pp. 1829–1838.e9.

Pawluk, A. et al. (2017) ‘Disabling a Type I-E CRISPR-Cas Nuclease with a Bacteriophage-Encoded Anti-CRISPR Protein’, mBio. doi: 10.1128/mbio.01751-17.

Potter, S. C., Luciani, A., Eddy, S. R., Park, Y., Lopez, R., & Finn, R. D. (2018). HMMER web server: 2018 update. Nucleic acids research, 46(W1), W200–W204.

Pul, U. et al. (2010) ‘Identification and characterization of E. coli CRISPR-cas promoters and their silencing by H-NS’, Molecular microbiology, 75(6), pp. 1495–1512.

Rauch, B. J. et al. (2017) ‘Inhibition of CRISPR-Cas9 with Bacteriophage Proteins’, Cell, 168(1-2), pp. 150–158.e10.

Richter, C. et al. (2012) ‘In vivo protein interactions and complex formation in the Pectobacterium atrosepticum subtype I-F CRISPR/Cas System’, PloS one, 7(12), p. e49549.

Rollie, C. et al. (2020) ‘Targeting of temperate phages drives loss of type I CRISPR-Cas systems’, Nature. doi: 10.1038/s41586-020-1936-2.

Rollins, M. F. et al. (2015) ‘Mechanism of foreign DNA recognition by a CRISPR RNA-guided surveillance complex from Pseudomonas aeruginosa’, Nucleic acids research, 43(4), pp. 2216–2222.

Rostøl, J. T. and Marraffini, L. (2019) ‘(Ph)ighting Phages: How Bacteria Resist Their Parasites’, Cell host & microbe, 25(2), pp. 184–194.

Serfiotis-Mitsa, D. et al. (2010) ‘The structure of the KlcA and ArdB proteins reveals a novel fold and antirestriction activity against Type I DNA restriction systems in vivo but not in vitro’, Nucleic acids research, 38(5), pp. 1723–1737.

Shehreen, S. et al. (2019) ‘Genome-wide correlation analysis suggests different roles of CRISPR-Cas systems in the acquisition of antibiotic resistance genes in diverse species’, Philosophical transactions of the Royal Society of London. Series B, Biological sciences, 374(1772), p. 20180384.

Shin, J. et al. (2017) ‘Disabling Cas9 by an anti-CRISPR DNA mimic’, Science Advances. doi: 10.1126/sciadv.1701620.

Shintani, M., Suzuki-Minakuchi, C. and Nojiri, H. (2015) ‘Nucleoid-associated proteins encoded on plasmids: Occurrence and mode of function’, Plasmid, 80, pp. 32–44.

Skennerton, C. T. et al. (2011) ‘Phage encoded H-NS: a potential achilles heel in the bacterial defence system’, PloS one, 6(5), p. e20095.

Söding, J., Biegert, A. and Lupas, A. N. (2005) ‘The HHpred interactive server for protein homology detection and structure prediction’, Nucleic acids research, 33(Web Server issue), pp. W244–8.

Song, G. et al. (2019) ‘AcrIIA5 Inhibits a Broad Range of Cas9 Orthologs by Preventing DNA Target Cleavage’, Cell reports, 29(9), pp. 2579–2589.e4.

Stanley, S. Y. et al. (2019) ‘Anti-CRISPR-Associated Proteins Are Crucial Repressors of Anti-CRISPR Transcription’, Cell, 178(6), pp. 1452–1464.e13.

Sullivan, M. J., Petty, N. K., & Beatson, S. A. (2011). Easyfig: a genome comparison visualizer. Bioinformatics (Oxford, England), 27(7), 1009–1010. https://doi.org/10.1093/bioinformatics/btr039

Thoma, S. and Schobert, M. (2009) ‘An improved Escherichia coli donor strain for diparental mating’, FEMS microbiology letters, 294(2), pp. 127–132.

Thomson, N. R. et al. (2000) ‘Biosynthesis of carbapenem antibiotic and prodigiosin pigment in Serratia is under quorum sensing control’, Molecular microbiology, 36(3), pp. 539–556.

Vercoe, R. B. et al. (2013) ‘Cytotoxic chromosomal targeting by CRISPR/Cas systems can reshape bacterial genomes and expel or remodel pathogenicity islands’, PLoS genetics, 9(4), p. e1003454.

Vial, L. and Hommais, F. (2019) ‘Plasmid-chromosome cross-talks’, Environmental Microbiology. doi: 10.1111/1462-2920.14880.

Watters, K. E. et al. (2018) ‘Systematic discovery of natural CRISPR-Cas12a inhibitors’, Science, 362(6411), pp. 236–239.

Watters, K. E. et al. (2020) ‘Potent CRISPR-Cas9 inhibitors from Staphylococcus genomes’, PNAS. doi: 10.1101/799403.

Westra, E. R. et al. (2010) ‘H-NS-mediated repression of CRISPR-based immunity in Escherichia coli K12 can be relieved by the transcription activator LeuO’, Molecular microbiology, 77(6), pp. 1380–1393.

Westra, E. R. et al. (2013) ‘Type I-E CRISPR-Cas Systems Discriminate Target from Non-Target DNA through Base Pairing-Independent PAM Recognition’, PLoS Genetics, p. e1003742. doi: 10.1371/journal.pgen.1003742.

Xu, Z. et al. (2019) ‘Native CRISPR-Cas-Mediated Genome Editing Enables Dissecting and Sensitizing Clinical Multidrug-Resistant P. aeruginosa’, Cell Reports, pp. 1707–1717.e3. doi: 10.1016/j.celrep.2019.10.006.

Yin, Y., Yang, B., & Entwistle, S. (2019). Bioinformatics Identification of Anti-CRISPR Loci by Using Homology, Guilt-by-Association, and CRISPR Self-Targeting Spacer Approaches. mSystems, 4(5), e00455–00419. doi:10.1128/mSystems.00455-19

Zavilgelsky, G. B. and Rastorguev, S. M. (2009) ‘Antirestriction proteins ArdA and Ocr as efficient inhibitors of type I restriction-modification enzymes’, Molecular Biology, pp. 241–248. doi: 10.1134/s0026893309020071.

Zhang et al. (2018) ‘CRISPRminer is a knowledge base for exploring CRISPR-Cas systems in microbe and phage interactions’, Communications Biology. doi: 10.1038/s42003-018-0184-6.

Zhang, H. et al. (2019) ‘Structural Basis for the Inhibition of CRISPR-Cas12a by Anti-CRISPR Proteins’, Cell host & microbe, 25(6), pp. 815–826.e4.

